# Effect of number of diffusion encoding directions in Neonatal Diffusion Tensor Imaging using Tract-Based Spatial Statistical analysis

**DOI:** 10.1101/2023.03.07.531625

**Authors:** Harri Merisaari, Linnea Karlsson, Noora M. Scheinin, Satu Shulist, John D. Lewis, Hasse Karlsson, Jetro J. Tuulari

**Affiliations:** FinnBrain Birth Cohort Study, Turku Brain and Mind Center, Department of Clinical Medicine, University of Turku, Turku, Finland; Department of Radiology, Turku University Central Hospital, Turku, Finland; Montreal Neurological Institute, McGill University, Montreal, Canada; Department of Medical Physics, Turku University Hospital, Turku, Finland; Department of Paediatrics and Adolescent Medicine, Turku University Central Hospital and University of Turku, Turku, Finland; Department of Psychiatry, Turku University Hospital and University of Turku; Centre for Population Health Research, Turku University Central Hospital and University of Turku, Turku, Finland; Turku Collegium of Science, Medicine and Technology, University of Turku, Turku, Finland

**Keywords:** Diffusion Tensor Imaging, Infant brain imaging, Tract-based spatial statistics

## Abstract

Diffusion Tensor Imaging (DTI) has been used to study the developing brain in early childhood, infants and *in utero* studies. In infants, number of used diffusion encoding directions has traditionally been smaller in earlier studies down to the minimum of 6 orthogonal directions. While the more recent studies often involve more directions, number of used directions remains an issue when acquisition time is optimized without compromising on data quality and in retrospective studies. Variability in the number of used directions may introduce bias and uncertainties to the DTI scalar estimates that affect cross-sectional and longitudinal study of the brain.

We analyzed DTI images of 133 neonates, each data having 54 directions after quality control, to evaluate the effect of number of diffusion weighting directions from 6 to 54 with interval of 6 to the DTI scalars with Tract-based spatial statistics (TBSS) analysis. The TBSS analysis was applied to DTI scalar maps, and the mean Region of Interest (ROI) values were extracted using JHU atlas.

We found significant bias in ROI mean values when only 6 directions were used (positive in FA, negative in MD, AD, RD), while when using 24 directions and above, the difference to scalar values calculated from 54 direction DTI was negligible.

Using DTI measurements from data with at least 24 directions may be used in comparisons with DTI measurements from data with higher numbers of directions.

## Introduction

Diffusion Tensor Imaging (DTI) is an established methodology for the study of the developing brain (Dubois et al., 2014; Lebel et al., 2019; Neil et al., 1998); it can be used to study typical brain development and in the study of the mechanisms of pathologies (Bui et al., 2006; Dubois et al., 2014; Dudink et al., 2007; McGraw et al., 2002; Miller et al., 2002; Neil et al., 2002; Partridge et al., 2004; Prayer and Prayer, 2003; Tanner et al., 2000). DTI has been used to study the developing brain in early childhood, in infancy (Forbes et al., 2002; Hüppi et al., 1996; Mukherjee et al., 2001; Neil et al., 1998), and in fetuses *in utero* (Bui et al., 2006; Jiang et al., 2009; Kasprian et al., 2008; Righini et al., 2003). Large changes occur in the first 24 months after birth in morphology and microstructure, which is tightly linked to rapid white matter myelination during this period of development (Mukherjee et al., 2001).

DTI studies have included pre-mature infant scans (Aeby et al., 2012, 2009; Miller et al., 2002; Neil et al., 1998; Partridge et al., 2004) and term born infants (Aeby et al., 2009; Forbes et al., 2002; Geng et al., 2012; Hermoye et al., 2006; McGraw et al., 2002; Mukherjee et al., 2001; Sadeghi et al., 2013). The commonly used DTI measures are Axial, Radial and Mean Diffusivities (AD, RD and MD) that all have a physical unit (mm^2^/s). Fractional Anisotropy (FA) is a degree of anisotropy of diffusion, and a proportional estimate of directionality. These DTI measures are expected to be in the same range when obtained from similar subjects in brain white matter, while naturally the scanner manufacturer and model, acquisition parameters, implemented quality control procedures, preprocessing and imaging data analysis method also likely affect the DTI scalar estimates. The used b-values (diffusion gradient strengths) have varied from 500 (Jiang et al., 2009) to 2600 s/mm^2^ (Bastiani et al., 2019). Number of used diffusion encoding directions in acquisition has generally been smaller in earlier studies down to minimum of 6 orthogonal directions, while the more recent studies have involved more directions (Batalle et al., 2019; Lebenberg et al., 2019). DTI data collected as a part of the developing Human Connectome Project (dHCP) represents the state-of-the-art in the field, using 88 encoding directions acquisition with b=1000 s/mm^2^. For details on the whole multi shell sequence please refer to (Bastiani et al., 2019), and to **Supplementary Information Table 1** and (Cachia et al., 2022) for literature review on infant brain DTI.

Earlier simulation study has suggested that as long as optimum encoding is used for six directions, no significant advantage is gained by adding more in terms of a quality measure normalized for imaging time (Hasan et al., 2001). In simulations with different acquisition protocols from 6 to 39 (Landman et al., 2007), bias in DTI scalars was noted, while suggesting that due to similar DTI contrasts the different procotols can be considered comparable in clinical studies as long as sampling orientations are balanced. However, in (Papadakis et al., 2000), directions from 18 to 21 were suggested as most efficient number of directions according to simulation experiments with various levels of Signal-to-Noise-Ratio (SNR). In addition to simulations, five healthy subjects were used in (Skare et al., 2000), demonstrating effectiveness of number of diffusion directions instead of signal averages for optimal FA measurement Further, in (Jones, 2004) it was suggested that 20 directions are required for proper estimation of FA, while for tensor orientation and mean diffusivity, 30 directions were suggested to avoid variations attributed to the orientation of the tissue. While the simulation studies provide good theoretical basis particularly for designing DTI acquisition protocol, they have not considered spatially heterogenous nature of brain *in vivo*, nor effects from different types of motion, occuring particularly during acquisition of infants.

For choosing between signal averaging and number of directions, it has been suggested to invest acquisition time into different diffusion directions (Correia et al., 2009). In tractography-based regions of interest analysis on DTI scalars evaluated in 13 healthy adults (Barrio-arranz et al., 2015), considerable changes were shown between 6, 21, 40 and 61 directions in the DTI scalars, with decrease in FA, AD and increase in RD when higher number of directions were used. Ten healthy adults were scanned with six acquisitions in two sessions for intra-session and inter-session variability in (Wang et al., 2012). DTI acquisitions with 15 and 30 directions was compared, and differences were found between them in nine evaluated fiber tracts. In another study, DTI-derived brain connectivity was analyzed in (Vaessen et al., 2010) with six healthy adults scanned twice in six gradient sampling schemes with number of gradient directions as 6, 15 and 32, and significant differences were found in small world brain topology metrics.

The effect of number of diffusion directions has been explored in adults. In one study with fifteen healthy adults (Ni et al., 2006) no significant difference was found between using 6, 21, and 31 directions, while number of excitations were, 10, 3, and 2 correspondingly. Similarly, (Widjaja et al., 2009) a study evaluated 7,15 and 25 directions with 10 healthy adults, finding no significant differences in FA. Ten major fiber tracts in eleven healthy adult volunteers was used to compare 6 direction data to 30 and 60 direction data, in (Lebel et al., 2011). Measures with the 6 directions data were considered similar to 30 and 60 directions, while also noting benefits of using higher number of directions. In contrast to studies supporting the use of only 6 directions for reliable DTI scalars, study with 6 adults in (Giannelli et al., 2010) used 6, 11, 19, 27 and 55 directions for measured contrast-to-signal variance between the main white matter and surrounding cerebral region. Use of more than 20 diffusion directions was suggested while keeping same number of directions within study, to avoid bias. In addition to changes in DTI scalars, (Barrio-arranz et al., 2015) reported changes also in tractography, in study with 13 healthy adults, comparing between 6, 21, 40 and 61 directions. In a multicenter study with two healthy adults and five imaging sites (Sakaie et al., 2012) there was high concordance between sites in FA an MD between when 33 encoding directions were used. Lastly, a study on data harmonization between two sites, with participants of 8 to 22 years old (Fortin et al., 2017), using two datasets having 105 subjects each, reported significant discrepancies in FA and MD between imaging sites having different hardware and different acquisition protocols, including diffusion directions of 30 and 64. Taken together, investigators have reached widely different conclusions for the degree to which the amount of diffusion-encoding directions is acceptable.

It may be tempting to use as few diffusion directions as possible (to the minimum of required 6 for DTI) as inclusion criteria for neurodevelopmental study for sake of shorter scanning time, and for allowing to keep as many cases as possible included the analysis, without needing them to be rejected due to too low number of directions after quality control procedures, e. g. due to corrupted or interrupted data acquisition (Dubois et al., 2006). Short term repeatability of paediatric brain DTI was studied in (Carlson et al., 2014) with two readers performing tracktography on 47 subjects with age of 4-18, and in (Harri Merisaari et al., 2019) for 122 neonates, giving results in reliability of measurements with the applied number of diffusion directions. However, not only precision and repeatability, but alo bias in the DTI scalars may cause serious problems, especially in longitudinal and neurodevelopmental studies, when subjects have different number of diffusion directions within and between different age groups (Tamnes et al., 2018). To the best of our knowledge, effect of number of DTI gradient directions has not been assessed in infants, which is crucial time point in neurodevelopmental studies, and earlier studies have generally focused only on FA and MD. Also, evaluations of DTI are needed to verify that in statistically significant findings in larger brain study cohorts acquired with different setting are not due to differences in their acquisition parameters.

To summarize, the earlier DTI infant literature has used various numbers of diffusion encoding directions, without assessing the effect of the number of directions. Simulation studies have made efforts to demonstrate the effect, but the results may be hard to apply more widely to real life situations for brain DTI acquired at varying age, particularly in infants where brain tissue characteristics and acquisition circumstances largely differ from those of adults. Thus, *in vivo* validation of simulation results (Jones, 2004) and earlier results in adults, is warranted for their applicability in infant DTI.

In the current study, we evaluated effect of number of diffusion weighting directions to the DTI scalars in context of Tract-based spatial statistics (TBSS) analysis, applied to all DTI scalars in 133 subjects. We used number of diffusion directions from 6 to 54 to systematically map the minimum amount of diffusion-encoding directions needed for reliable tensor estimates (FA, MD, AD and RD).

## Materials and Methods

The study was conducted according to the Declaration of Helsinki and was reviewed and approved by the Ethics Committee of the Hospital District of Southwest Finland (ETMK: 31/180/2011). The imaging data that support the findings of this study are available upon reasonable request from the corresponding author, while the imaging data are not publicly available due to privacy restrictions.

### Subjects

The participants were from the FinnBrain Birth Cohort study (Karlsson et al., 2018), which enrolled families to a neonatal MRI study (Lehtola et al., 2020). The families were recruited at three healthcare locations in Southwest Finland during their first trimester ultrasound visit at gestational week (GW) 12. From this broader participant pool, 189 newborn-mother dyads were recruited into this study. They were recruited based on willingness to participate, availability of the newborn to have an MRI two to five weeks after birth, childbirth being after GW 31, birthweight more than 1500 g, and not having a previously diagnosed central nervous system (CNS) anomaly or abnormal findings in a previous MRI scan. After explaining the study’s purpose and protocol, written informed consent was obtained from the parent(s). Of these 189 newborn participants, 54 had motion artefacts in the MR images. Additionally, two mothers had missing prenatal distress questionnaires. Therefore 133 newborn-mother dyads were eligible for statistical analyses. For the current study, we included all neonates that had at least 54 successful unique diffusion encoding directions in their DTI data following rigorous quality control outlined below, see **Table 1** for demographics of the study.

**Table 1.** Subject demographics.

|  |  |  |
| --- | --- | --- |
| **Infant sex** | **N** | **Percent** |
| **male** | **69** | **51.8%** |
| **female** | **64** | **48.2%** |
| **Infant age in weeks** | **Mean** | **SD** |
| **Gestational age (from estimated conception)** | **43.6** | **1.04** |
| **Age from term age (from gw 40)** | **3.5** | **1.04** |
| **Age from birth to scan** | **3.7** | **1.10** |
| **Birth characteristics** | **Mean** | **SD** |
| **Birth weight (g)** | **3487** | **436** |
| **Birth height (cm)** | **50.4** | **1.86** |
| **Mother characteristics** | **Mean** | **SD** |
| **Maternal age at birth (years)** | **29.6** | **4.4** |
| **Maternal body mass index (BMI)** | **24.2** | **4.0** |
| **Educational level (low / middle / high)** | **37 / 43 / 49 [4 missing]** |  |
| **Maternal smoking during pregnancy (yes / no)** | **11 / 122** |  |
| **SSRI / SNRI medication in use (yes / no)** | **12 / 121** |  |

### Image Acquisition

In the DTI scans, a 12-element Head Matrix coil was used in a twice-refocused Spin Echo-Echo Planar Imaging (SE-EPI) sequence. A diffusion weighting of b=1000 s/mm^2^ was used with 2×2×2 mm^3^ isotropic resolution (FOV 208 mm, 64 slices, TR/TE=8500/90 ms) the data was acquired in three segments composing of a total 96 uniformly distributed directions. The DTI acquisition scheme is provided in detail in (H. Merisaari et al., 2019).

### Image Preprocessing and TBSS analysis

Data were pre-processed as described in (H. Merisaari et al., 2019). In brief, the images were first quality controlled with DTIprep (Oguz et al., 2014), and motion corrupted volumes were removed from each segment. Fieldmap correction was not applied here as we did not have opposing phase directions in the sequences (distortions were minimal for the current data). FSL tools were used to derive the tensor maps (FA, MD, AD and RD). For the purposes of the current study, we created a sub-set from the whole sample so that included cases had at least 54 / 96 acceptable directions after quality control. The dataset had 133 / 189 participants. We chose 54 directions as threshold limit to be included, as it was considered a suitable trade-off between number of subjects to be included, and number of directions for evaluations. From dataset with 54 directions, individual directions were dropped out while keeping diffusion direction distribution as uniform as possible by maximizing the angular resolution using algorithm described in (H. Merisaari et al., 2019), to create simulated versions of increasing diffusion-encoding directions: 6,12,18, 24, 30, 36,42,48 and 54 directional data sets.

TBSS analysis (Smith et al., 2006) was first applied to each datasets containing 54 directions with FA values, and the mean FA image was then used as a template for all data sets. This generated spatially aligned images of DTI scalars for evaluations of number of diffusion encoding directions in voxels of DTI parameter maps aligned to the TBSS template. In addition to voxel-level analysis, to support the ROI analyses, John Hopkins University DTI-based white matter atlas for neonates (Oishi et al., 2011) (JHU atlas) was applied to the TBSS processed DTI scalar maps. The mean intensity values in 16 most important Regions of Interest (ROI) were then used bilaterally to analyze differences between number of used diffusion directions.

### Statistical Analysis

*We* evaluated difference of using direction at voxel level using repeated measures analysis followed by paired t-test as post hoc test (8 tests comparing 6, 12, 18, 24, 30, 36, 42, 48 directions against 54 directions), using fsl toolbox randomise tool (Winkler et al., 2014). As the scalar value distributions in ROIs were found deviating from normal distribution, we evaluated ROI mean value differences with Friedman test and Wilcoxon signed rank post hoc test. In DTI, smaller number of directions are expected to affect FA bias towards higher values (Pierpaoli and Basser, 1996). Thus, repeated measures Mann-Kendall test was used to test the existence of a non-zero trend in change of DTI scalar ROI values across number of directions. ICC(2,1) was applied between directions, measuring absolute agreement with scalar measures, to see their suitability for studies comparing patient groups with controls, or longitudinal studies. We considered ICC(2,1) > 0.75 to be excellent, while < 0.75 to be poor or moderate (Koo and Li, 2016). All statistics were performed with R version 3.6.3 (Team, 2020). P-values below 0.05 after False Discovery Rate (FDR) correction for multiple comparisons were considered statistically significant, unless otherwise noted.

## Results

The DTI scalar values from images with 54 directions were used as benchmark, i.e. the best possible estimate of the true tensor values in the analyses. The reference fractional anisotropy values across TBSS skeleton are shown in **Figure 1** to visualize the typical values observed in our data (see **Supplementary Information Figure 1** for corresponding values in MD, AD, RD), where high FA values are found in the midsagittal callosum and central regions in superior fasciculus, while lower FA values are located in peripheral regions.

**Figure 1.**
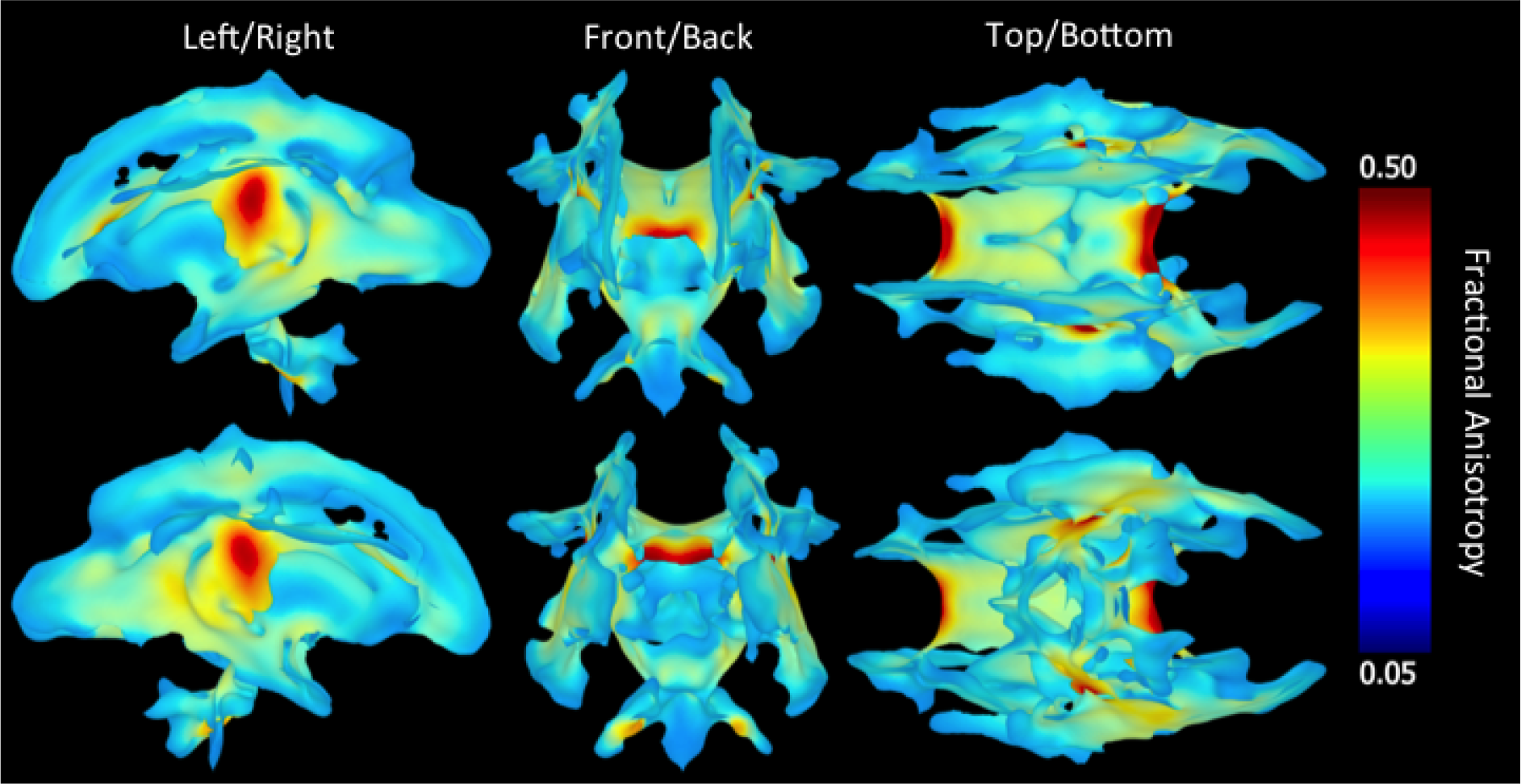
Rendering of DTI Fractional Anisotropy map of Tract Based Spatial Statistics (TBSS) map in six orthogonal directions, calculated using 54 diffusion encoding directions in 133 neonates.

We calculated how much minimum requirement for number of diffusion from 54 to 96 directions after post-processing, would affect to the number of included subjects. The results are shown in **Figure 2.** As the criteria for available directions is increased from 55 to the maximum of 96, we observed generally a linear descending trend in the number of cases, with minor change at 80 directions.

**Figure 2.**
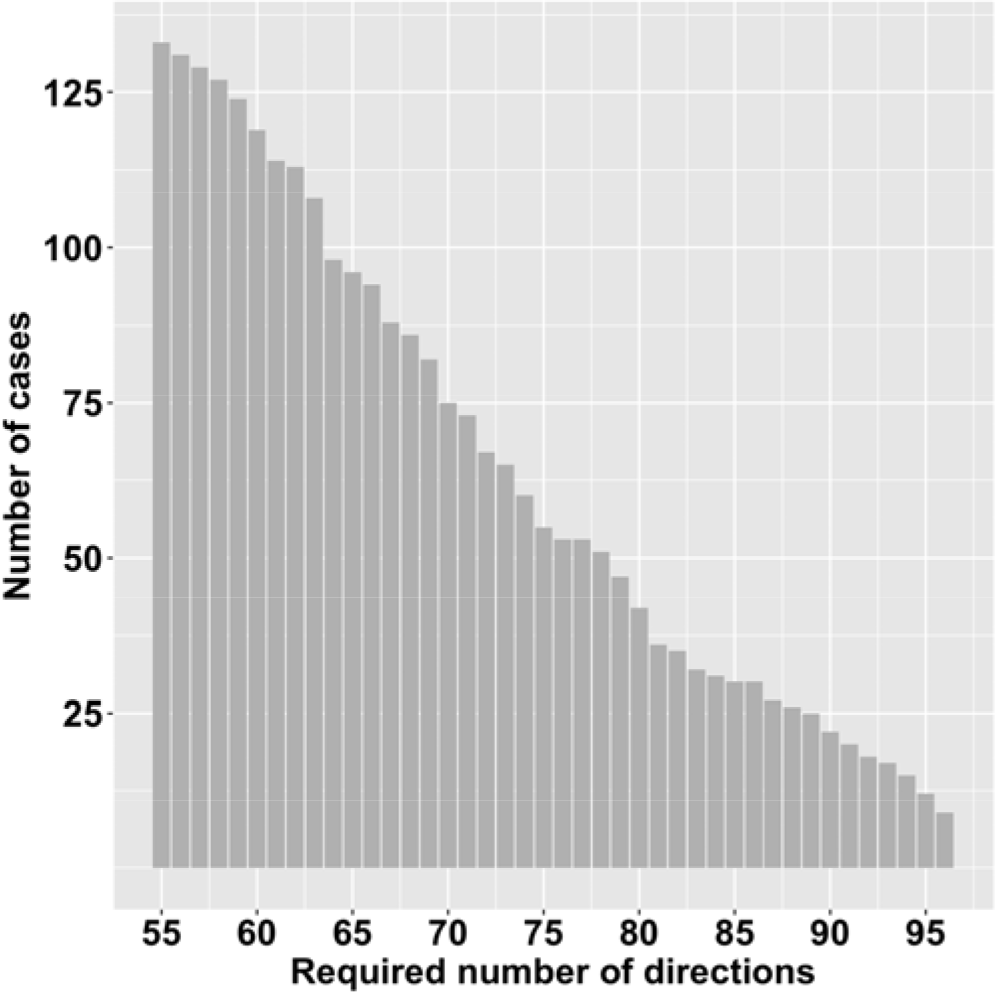
Effect of raising exclusion criteria for DTI images starting from 55 directions where 133 subjects were available, and ending to the maximum of 96 directions used in the full acquisition.

### Repeated measures voxel-wise analysis

In voxelwise repeated measures analysis, all TBSS skeleton locations were found to differ in individuals statistically significantly (p<0.001 after correction for multiple comparisons across TBSS voxels) in comparison to 54 direction scalar estimates. The corresponding absolute scalar value differences were evaluated for TBSS voxels by visualizing percentage of subjects where a DTI scalar value was 10% or more different than corresponding value in 54 direction data. The results for FA for 6 to 48 directions, are shown in **Figure 3** (see **Supplementary Information Figure 2** for MD, AD, RD correspondingly). With number of 6, 12 and 18 directions, large regions across TBSS voxels demonstrated significant proportions (~10% or more) of subjects were deviating more than 10% from the reference values.

**Figure 3.**
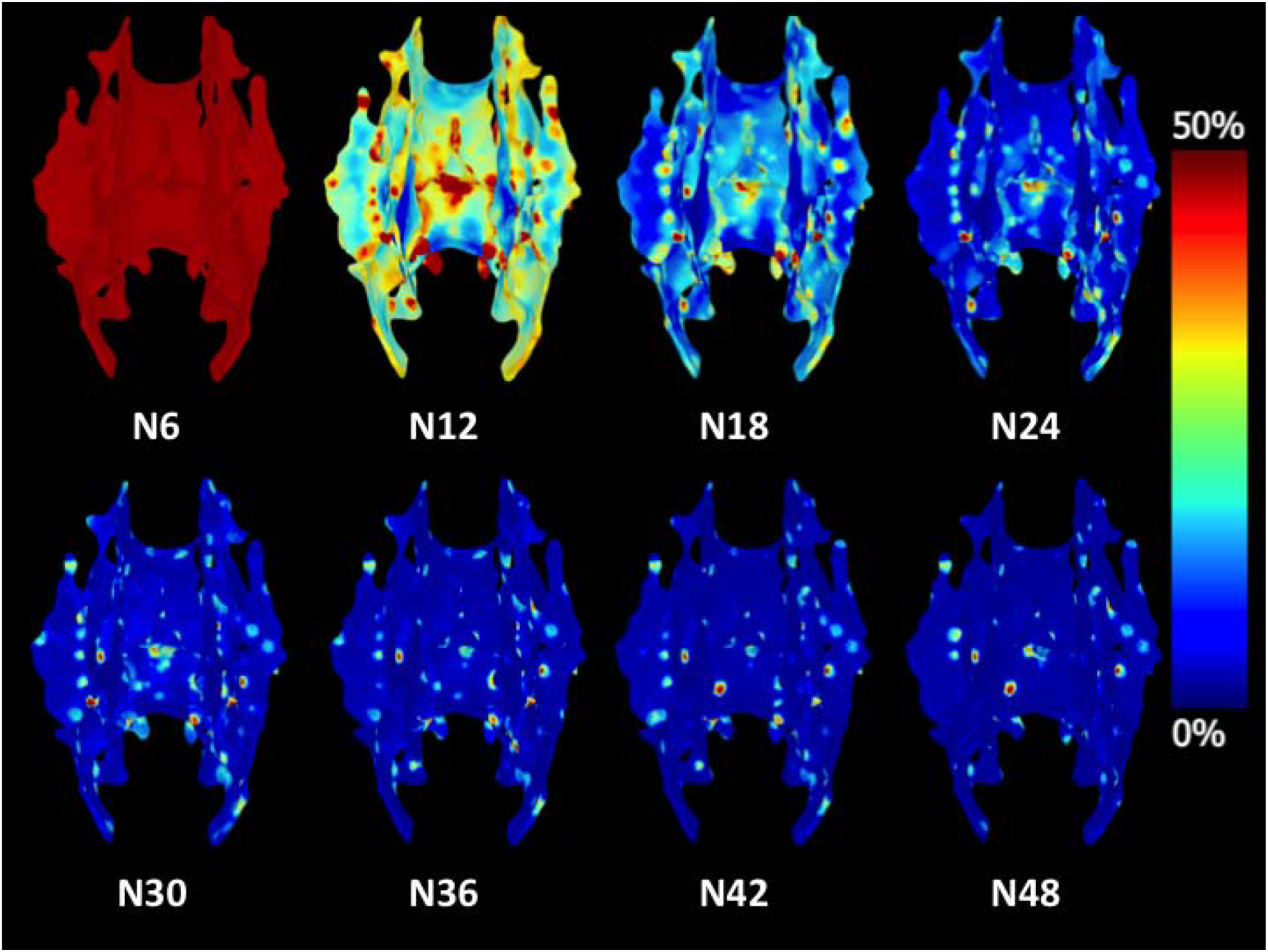
Rendering of proportions of neonatal subjects (%) from 133, where difference in Fractional Anisotropy was larger than 10%, when comparing values from 6(N6) to 48(N48) diffusion encoding directions to values with 54 directions. Directions 6, 12 and 18 contain larger regions where non-negligible proportion of subjects deviate more than 10% from the reference value with 54 directions.

### Repeated measures ROI-wise analysis

When comparing ROI mean values in TBSS skeleton, for MD, AD and RD, only 6 direction data was significantly different from 54 direction values (p<0.05). The differences were within all of the 16 evaluated regions (see **Supplementary information figure 3** for list of abbreviations) at either side, apart from Cingulum Cingular Part. In FA, the 12 direction values were significantly different in 12 regions at either side (p<0.05), the 18 direction values we re different in 6 regions (p<0.05), and 24 direction values in 3 regions (p<0.05 for Cerebral peduncle, Corticospinal tract and Cingulum cingular part). Of those three, Cingulum cingular part (right) was also significantly different in all directions up to 42 directions.

### Effect of angular resolution to the ROI-level results

*We* also evaluated Spearman correlation of mean angular resolution and number of directions, and evaluated interaction of directions and angular resolution in explaining standard deviation of scalar values across subjects. The angular resolution was found to interact with number of directions in all of the evaluated ROIs (p<0.05). Spearman correlation between angular resolution and scalar value was found significant with 6 directions (p<0.05), while no statistically significant correlation was found in other numbers of directions from 12 to 54. The variance of the scalar value was particularly high with 6 direction data (see **Supplementary Information Figure 3** for box plots with 6, 24, 54 directions). Based on found voxel-wise and ROI-wise differences, we excluded directions 6,12,18 from subsequent analysis, to focus on examining if having larger number of directions 24 to 54 would be considered interchangeable.

### Mann-Kendall difference between diffusion encoding directions from 24 to 54 in ROI values

With Mann-Kendall test for monotonic trend in DTI scalars when number of diffusion directions varied from 24 to 54, we found statistically significant positive trend after FDR correction (p<0.05) in FA of Corpus Callosum (CC), Retrolenticular internal capsule (RLIC) and External capsule (EC). However, median FA difference between 24 and 54 directions was considered negligible (FA difference < 0.01). In AD, significant negative trend was found in Cingululum cingular part (CGC) with median difference of −0.3×10^-3^mm^2^/s (left), and −0.2×10^-3^ mm^2^/s (right), between 24 and 54 directions. In MD, significant negative trends were found in CGC (−0.2×10^-3^ mm^2^/s left and right) and Cerebral peduncle (−0.1×10^-3^ mm^2^/s). In RD, significant negative trend was found in CGC (−0.2×10^-3^ mm^2^/s left, 0.1×10^-3^ mm^2^/s right) and Superior fronto-occipital fasciculus (0.1×10^-3^ mm^2^/s right). Other trends in AD, MD and RD were either not significant (FR corrected p>0.05) or the scalar value difference between 24 and 54 was considered negligible (median difference <0.01×10^-3^mm^2^/s).

### Comparison of 24 and 54 diffusion encoding directions in voxel-wise analysis

We calculated also voxelwise differences across the TBSS skeleton are shown in **Figure 4** for absolute differences in FA between 24 and 54 direction data (see **Supplementary Information Figure 4** for MD, AD and RD, correspondingly), as these measures can be considered as upper limit for scalar value differences for 24 directions and higher. Generally, there were individual voxels demonstrating higher dependence on the number of directions, while the majority of the TBSS skeleton voxels had negligible differences between using 24 and 54 diffusion directions.

**Figure 4.**
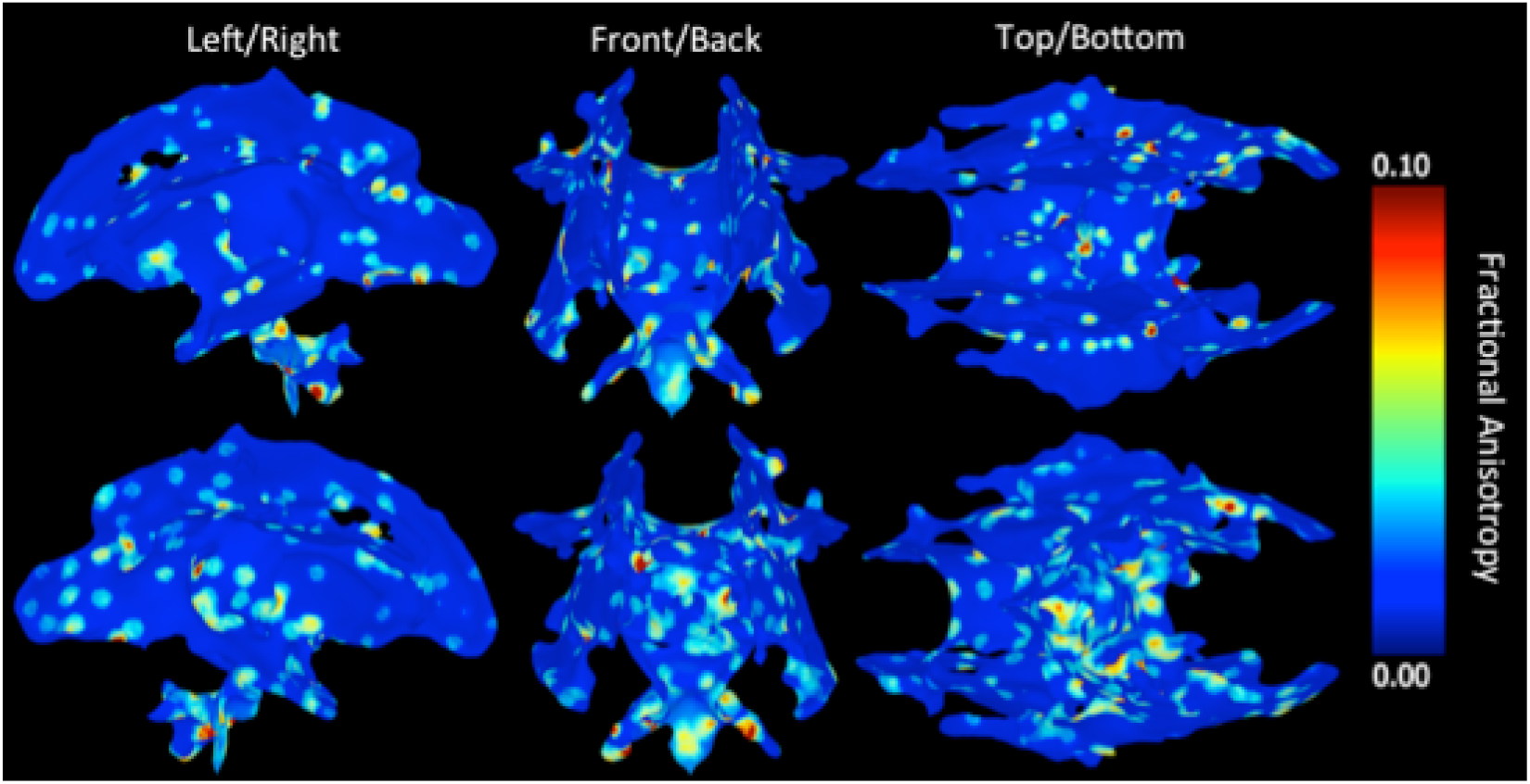
Rendering of Fractional Anisotropy (FA) difference, calculated using difference between using 24 and 54 encoding directions, in 133 neonates.

### *Voxel-wise ICC(2,1) for 24*, 30, 36,42, 48 *and 54 diffusion encoding directions*

ICC(2,1) reflects to absolute agreement in the scalar values when number of directions is considered as measurement methods, high ICC(2,1) indicating that the measurement are reproducible when directions are varied from 24 to 54. **Figure 5** demonstrates voxel-wise mapping of the ICC values across TBSS skeleton for FA (see **Supplementary Information Figure 5** for MD, AD and RD). In FA, ICC(2,1) (>0.75) was good across TBSS skeleton. Correspondingly, in AD, RD and MD, ICC(2,1) was generally lower, while central regions still expressing generally good reproducibility (ICC>0.75). Similar to measurements of absolute differences between 24 and 54 directions, individual locations with lower ICC(2,1) reproducibility were found to be present across TBSS skeleton, while majority of the skeleton had high reproducibility.

**Figure 5.**
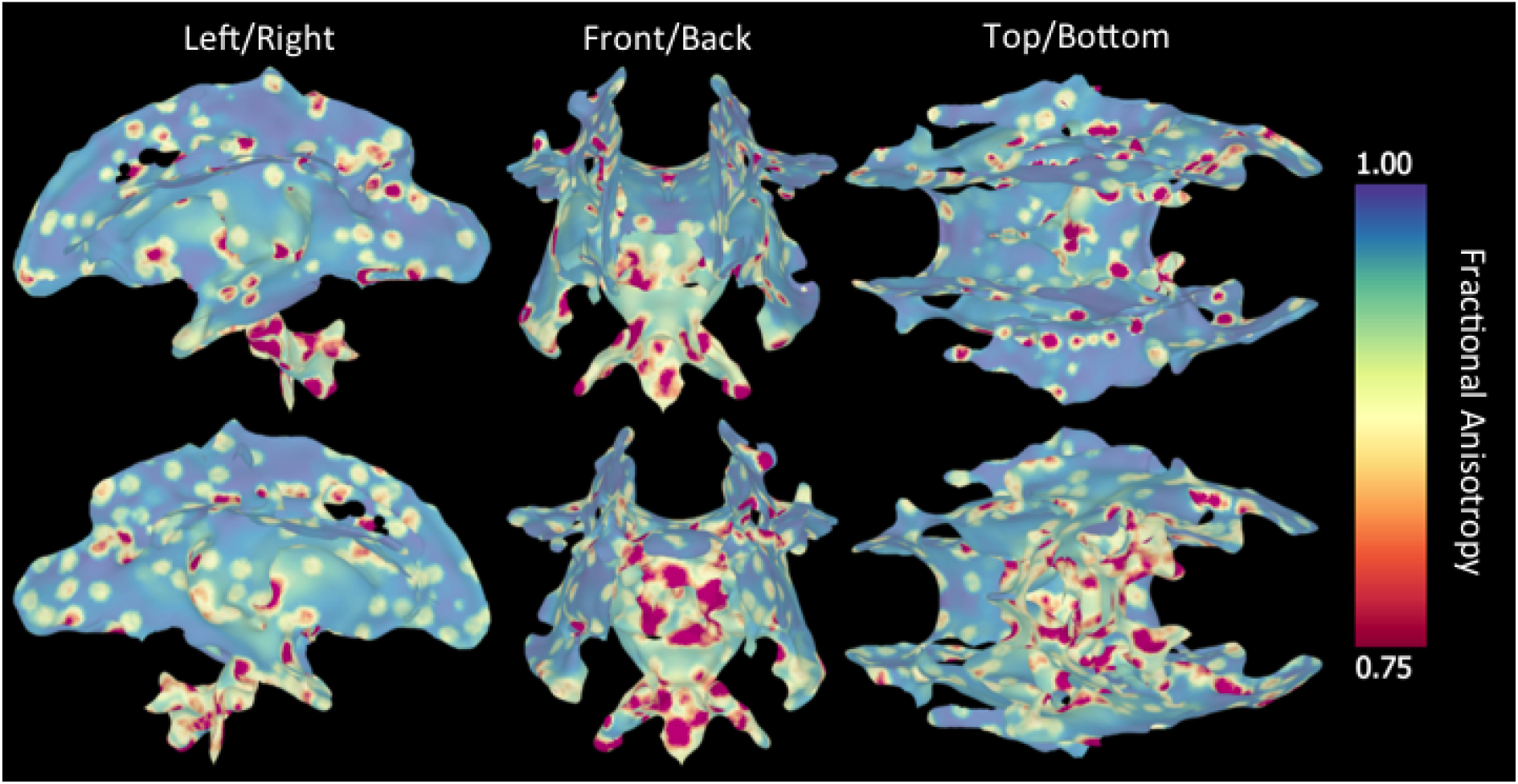
Rendering of Infraclass Correlation Coeffcient (ICC(2,1)) map of Fractional Anisotropy values, calculated using 24, 30, 36, 42, 48 and 54 encoding directions, in 133 neonates. Note that the colour coding starts from ICC 0.75, which is commonly considered sufficient threshold to pass reliability testing.

## Discussion

Various diffusion encoding strategies have been applied in studies of neurodevelopment at the microstructural level in neonates, but effect of diffusion directions has not been evaluated *in vivo* in larger extent before. The DTI scalar values from acquisitions at early age form the basis for further longitudinal differential comparisons, and therefore the accuracy of the scalar measurements is of high importance. We evaluated the effect of the number of diffusion encoding directions on the DTI scalar values in 2-5 weeks old subjects, where our analysis pipeline included quality control, motion and eddy correction, and final DTI scalar estimates extracted from TBSS scalar. We uniformized the 133 subjects to have the same number of diffusion directions for evaluations between directions from 6 to 54, in steps of 6 to observe the effect of the number of directions in that range. Even though the distribution of the angular resolution in the analysed data was minimized, our *in vivo* infant data had rotated b-vectors after motion correction. In contrast to studies where motion artefacts and thus related motion induced deviations from uniform diffusion direction distribution are not assessed, we consider our approach to provide more realistic evaluations. While this may have partially affected deviance of 6 direction data (as angular resolution correlated significantly with the scalar values) to 54 direction reference values, in 12 and 18 direction data we found no significant effect with angular resolution, and therefore we consider that their difference to 54 data is majorly due to number of directions rather than sensitivity to motion artefact induced changes from original acquired diffusion encoding directions. Our evaluations we mainly with TBSS skeleton values, as this technique can be considered robust against degradations in image quality. In addition to voxel-level analysis, we applied common approach of ROI-level analysis. As the sizes and shapes of ROIs make challenges for comparing between neurodevelopmental studies (Dudink et al., 2007), we applied automatic ROI placement in TBSS parameter maps to provide results in an easily applicable and reproducible context.

As expected, when exclusion criteria for number of diffusion directions was increased, the available number of subject reduced correspondingly (see **Figure 2).** This trade-off between number of directions and included cases is to be interpreted in context where choice for required minimum number of directions may depend on number of factors, such as size of the smallest or smaller compared study group, general inter-subject variance in DTI measure of interest, and general willingness to exclude obtained data in exchange of quality of included data.

The above in mind, we can generally consider that decision resulting in excluding more than 50% of subjects would have too low cost-benefit ratio for practical implementations. This would mean that requiring to have more than 70 directions with acceptable quality from original acquired 96, may be unrealistic with the type of cases used in this study. In theory, more acquisition time could be invested to obtain more original directions, and further, more acceptable quality data, given that challenges relating to neonatal image acquisition can be addressed for longer durations. In our experiments, for obtaining 24 quality-controlled directions, we estimate that at least close to 50 directions would be needed to be acquired during scan to compensate for loss of data due to expected artefacts. Naturally, the DTI acquisition success rate affects this estimate (in our study approximately 26% of cases were excluded), and imputation and other procedures may be used to replace corrupted or missing data.

*We* observed high systematic bias between using six directions and 54 directions confirming findings in theoretical and simulation studies (Jones, 2004), and studies with healthy adults relating to effect of the number of diffusion directions. In contrast to (Jones, 2004), we had motion induced effects *in vivo*, while onsidering also that the loss of directions may need to be compensated by acquiring extra directions to ensure desired amount in the final analysis after quality control. Our results suggest that having 24 directions and above should be considered suitable for purposes of deriving accurate DTI scalars in infant TBSS analyses, with either non-significant or negligible differences compared to using 54 directions. These results are largely in alignment also with the earlier studies with adults (Barrio-arranz et al., 2015; Giannelli et al., 2010) for which our results can be considered as confirmatory with larger *in vivo* sample, although with children, being in contrast to (Lebel et al., 2011).

The DTI scalar estimates were not largely found to change in higher number of directions from 24 up to 48 in comparison to 54-direction data (see **Figure 3,** and **Supplementary Information Figure 3),** with most of the subjects having less than 10% difference across the TBSS skeleton (see **Figure 3** and **Supplementary Information Figure 4).** This suggests that a dataset with varying number of diffusion directions between subjects could tentatively be combined into the same pooled neurodevelopmental sample, if number of diffusion directions is at least 24 in all of them, although more directions (Sakaie et al., 2012) and proper harmonization may be needed (Fortin et al., 2017). Larger numbers of diffusion directions are more challenging to acquire in infants / small children. Our results indicate that composing together DTI data having less than 24 directions with other DTI data with more directions may be problematic. We suggest that interpreting scalar estimates from data where smaller than 18 directions are available after quality control steps is discouraged due to potential bias introduced from small number of directions. However, this bias could naturally be reduced with producing uniform data with the caveat of losing acquired diffusion direction images, and thus accuracy of DTI measures.

While we simulated the varying amount of diffusion encoding directions based on single acquisition with 96 directions with consecutive quality control procedures, we expect the results to be the same if the actual image acquisition were performed completely separately for each of the number of direction from 6 to 54. This remains to be verified in future studies. TBSS is frequently used in similar settings, and we did not extend the studies on comparing possible enhancements with alternative spatial co-registration tools. Finally, it is left for future research to study the effect of diffusion acquisition analysis schemes, e.g. to tractography and after harmonization procedures.

## Conclusion

The number of diffusion directions leads to systematic bias in the DTI scalar measures, and using less than 24 diffusion directions is discouraged for neurodevelopmental studies. In evaluations of different numbers of diffusion encoding directions from 6 to 54, we observed that 24 quality-controlled directions in TBSS analysis provides negligible bias.

## Acknowledgements

We thank all the participating families, the staff of the Medical Imaging Centre of Turku University Hospital as well as the FinnBrain Birth Cohort Study research personnel. This research has received funding from the Jane and Aatos Erkko Foundation (HK), Academy of Finland (HM, #26080983), Hospital District of Southwest Finland State Research Grants (HM, JJT, NMS, HK), Alfred Kordelin Foundation, Turku University Foundation (JJT), Emil Aaltonen Foundation (JJT), and The Gyllenberg Foundation (NMS).

## References

Aeby, A., Bogaert, P. Van, David, P., Balériaux, D., Vermeylen, D., Metens, T., Tiège, X. De, 2012. Nonlinear microstructural changes in the right superior temporal sulcus and lateral occipitotemporal gyrus between 35 and 43 weeks in the preterm brain. Neuroimage 63, 104–110. https://doi.org/10.1016/j.neuroimage.2012.06.013

Aeby, A., Liu, Y., David, P., Bale, D., Metens, T., Bogaert, B. Van, 2009. Maturation of Thalamic Radiations between 34 and 41 Weeks ‘ Gestation□: A Combined Voxel-Based Study and Probabilistic Tractography with Diffusion Tensor Imaging. Am. J. Neuroradiol. 30, 1780–1786. https://doi.org/10.3174/ajnr.A1660

Barrio-arranz, G., Luis-garcía, R. De, Tristán-vega, A., Martín-, M., 2015. Impact of MR Acquisition Parameters on DTI Scalar Indexes□: A Tractography Based Approach. PLoS One 10, 1–19. https://doi.org/10.1371/journal.pone.0137905

Bastiani, M., Andersson, J.L.R., Cordero-Grande, L., Murgasova, M., Hutter, J., Price, A.N., Makropoulos, A., Fitzgibbon, S.P., Hughes, E., Rueckert, D., Victor, S., Rutherford, M., Edwards, A.D., Smith, S.M., Tournier, J.D., Hajnal, J.V., Jbabdi, S., Sotiropoulos, S.N., 2019. Automated processing pipeline for neonatal diffusion MRI in the developing Human Connectome Project. Neuroimage 185, 750–763. https://doi.org/10.1016/j.neuroimage.2018.05.064

Batalle, D., O’Muircheartaigh, J., Makropoulos, A., Kelly, C.J., Dimitrova, R., Hughes, E.J., Hajnal, J.V., Zhang, H., Alexander, D.C., Edwards, A.D., Counsell, S.J., 2019. Different patterns of cortical maturation before and after 38 weeks gestational age demonstrated by diffusion MRI in vivo. Neuroimage 185, 764–775. https://doi.org/10.1016/j.neuroimage.2018.05.046

Bui, T., François, J.D., Zaccaria, I., Alberti, C., Elmaleh, M., Garel, C., Luton, D., Blanc, N., Sebag, G., 2006. Microstructural development of human brain assessed in utero by diffusion tensor imaging. Pediatr. Radiol. 36, 1133–1140. https://doi.org/10.1007/s00247-006-0266-3

Cachia, A., Mangin, J., Dubois, J., 2022. Mapping the Human Brain from the Prenatal Period to Infancy Using 3D MAgnetic Reonance Imaging, in: Houdé, O., Borst, G. (Eds.), The Cambridge Handbook of Congitive Development. Cambridge Univerity Press, p. 50.

Carlson, H.L., Laliberté, C., Brooks, B.L., Hodge, J., Kirton, A., Bello-espinosa, L., Hader, W., Sherman, E.M.S., 2014. Epilepsy & Behavior Reliability and variability of diffusion tensor imaging (DTI) tractography in pediatric epilepsy. Epilepsy Behav. 37, 116–122. https://doi.org/10.1016/j.yebeh.2014.06.020

Correia, M.M., Carpenter, T.A., Williams, G.B., 2009. Looking for the optimal DTI acquisition scheme given a maximum scan time□: are more b-values a waste of time□? Magn. Reson. Imaging 27, 163–175. https://doi.org/10.1016/j.mri.2008.06.011

Dubois, J., Kulikova, S., Poupon, C., Hu, P.S., 2014. The early development of brain white matter: a review of imaging studies in fetuses, newborns and infants. Neuroscience 276, 48–71. https://doi.org/10.1016/j.neuroscience.2013.12.044

Dubois, J., Poupon, C., Lethimonnier, F., Le Bihan, D., 2006. Optimized diffusion gradient orientation schemes for corrupted clinical DTI data sets. Magn. Reson. Mater. Physics, Biol. Med. 19, 134–143. https://doi.org/10.1007/s10334-006-0036-0

Dudink, J., Lequin, M., Pul, C. Van, Buijs, J., Conneman, N., Goudoever, J. Van, Govaert, P., 2007. Fractional anisotropy in white matter tracts of very-low-birth-weight infants. Pediatr. Radiol. 37, 1216–1223. https://doi.org/10.1007/s00247-007-0626-7

Forbes, K.P., Pipe, J.G., Bird, C.R., 2002. Changes in Brain Water Diffusion during the 1st Year of Life. Radiology 222, 405–409.

Fortin, J., Parker, D., Tunç, B., Watanabe, T., Elliott, M.A., Ruparel, K., Roalf, D.R., Satterthwaite, T.D., Gur, R.C., Gur, R.E., Schultz, R.T., Verma, R., Shinohara, R.T., 2017. NeuroImage Harmonization of multi-site diffusion tensor imaging data. Neuroimage 161, 149–170. https://doi.org/10.1016/j.neuroimage.2017.08.047

Geng, X., Gouttard, S., Sharma, A., Gu, H., Styner, M., Lin, W., Gerig, G., Gilmore, J.H., Gerig, G., Gilmore, J.H., 2012. Quantitative tract-based white matter development from birth to age 2 years. Neuroimage 61, 542–557. https://doi.org/10.1016/j.neuroimage.2012.03.057

Giannelli, M., Cosottini, M., Michelassi, M.C., Lazzarotti, G., Belmonte, G., Bartolozzi, C., Lazzeri, M., 2010. Dependence of brain DTI maps of fractional anisotropy and mean diffusivity on the number of diffusion weighting directions. J. Appl. Clin. Med. Phys. 11, 176–190.

Hasan, K.M., Parker, D.L., Alexander, A.L., 2001. Comparison of gradient encoding schemes for diffusion-tensor MRI. J. Magn. Reson. Imaging 13, 769–780. https://doi.org/10.1002/jmri.1107

Hermoye, L., Saint-martin, C., Cosnard, G., Lee, S., Kim, J., Nassogne, M., Menten, R., Clapuyt, P., Donohue, P.K., Hua, K., Wakana, S., Jiang, H., Zijl, P.C.M. Van, Mori, S., 2006. Pediatric diffusion tensor imaging□: Normal database and observation of the white matter maturation in early childhood. Neuroimage 29, 493–504. https://doi.org/10.1016/j.neuroimage.2005.08.017

Hüppi, P.S., Schuknecht, B., Boesch, C., Bossi, E., Felblinger, J., Fusch, C., Herschkowitz, N., 1996. Structural and Neurobehavioral Delay in Postnatal Brain Development of Preterm Infants. Pediatr. Res. 39, 895–901.

Jiang, S., Xue, H., Counsell, S., Anjari, M., Allsop, J., Rutherford, M., Rueckert, D., Hajnal, J.V., 2009. Diffusion Tensor Imaging (DTI) of the Brain in Moving Subjects□: Application to In-Utero Fetal and Ex-Utero Studies. Magn. Reson. Med. 62, 645–655. https://doi.org/10.1002/mrm.22032

Jones, D.K., 2004. The Effect of Gradient Sampling Schemes on Measures Derived from Diffusion Tensor MRI: A Monte Carlo Study. Magn. Reson. Med. 51, 807–815. https://doi.org/10.1002/mrm.20033

Karlsson, L., Tolvanen, M., Scheinin, N.M., Uusitupa, H.M., Korja, R., Ekholm, E., Tuulari, J.J., Pajulo, M., Huotilainen, M., Paunio, T., Karlsson, H., 2018. Cohort Profile: The FinnBrain Birth Cohort Study (FinnBrain). Int. J. Epidemiol. 47, 15–16j. https://doi.org/10.1093/ije/dyx173

Kasprian, G., Brugger, P.C., Weber, M., Krssák, M., Krampl, E., Herold, C., Prayer, D., 2008. NeuroImage In utero tractography of fetal white matter development. Neuroimage 43, 213–224. https://doi.org/10.1016/j.neuroimage.2008.07.026

Koo, T.K., Li, M.Y., 2016. A Guideline of Selecting and Reporting Intraclass Correlation Coefficients for Reliability Research. J. Chiropr. Med. 15, 155–163. https://doi.org/10.1016/j.jcm.2016.02.012

Landman, B.A., Farrell, J.A.D., Jones, C.K., Smith, S.A., Prince, J.L., Mori, S., 2007. Effects of diffusion weighting schemes on the reproducibility of DTI-derived fractional anisotropy, mean diffusivity, and principal eigenvector measurements at 1.5T. Neuroimage 36, 1123–1138. https://doi.org/10.1016/j.neuroimage.2007.02.056

Lebel, C., Benner, T., Beaulieu, C., 2011. Six Is Enough? Comparison of Diffusion Parameters Measured Using Six or More Diffusion-Encoding Gradient Directions With Deterministic Tractography. Magn. Reson. Med. 68, 474–483. https://doi.org/10.1002/mrm.23254

Lebel, C., Treit, S., Beaulieu, C., 2019. A review of diffusion MRI of typical white matter development from early childhood to young adulthood. NMR Biomed. 32, e3778. https://doi.org/10.1002/nbm.3778

Lebenberg, J., Mangin, J.F., Thirion, B., Poupon, C., Hertz-Pannier, L., Leroy, F., Adibpour, P., Dehaene-Lambertz, G., Dubois, J., 2019. Mapping the asynchrony of cortical maturation in the infant brain: A MRI multi-parametric clustering approach. Neuroimage 185, 641–653. https://doi.org/10.1016/j.neuroimage.2018.07.022

Lehtola, S.J., Tuulari, J.J., Scheinin, N.M., Karlsson, L., Parkkola, R., Merisaari, H., Lewis, J.D., Fonov, V.S., Louis Collins, D., Evans, A., Saunavaara, J., Hashempour, N., Lähdesmäki, T., Acosta, H., Karlsson, H., 2020. Newborn amygdalar volumes are associated with maternal prenatal psychological distress in a sex-dependent way. NeuroImage Clin. 28, 102380. https://doi.org/10.1016/j.nicl.2020.102380

McGraw, P., Liang, L., Provenzale, J.M., 2002. Evaluation of Normal Age-Related Changes in Anisotropy During Infancy and Childhood as Shown by Diffusion Tensor Imaging. Am. J. Roentgenol. 179, 1515–1522.

Merisaari, Harri, Tuulari, J.J., Karlsson, L., Scheinin, N.M., Parkkola, R., Saunavaara, J., Lähdesmäki, T., Lehtola, S.J., Keskinen, M., Lewis, J.D., Evans, A.C., Karlsson, H., 2019. Test-retest reliability of Diffusion Tensor Imaging metrics in neonates. Neuroimage 197, 598–607. https://doi.org/10.1016/j.neuroimage.2019.04.067

Merisaari, H., Tuulari, J.J., Karlsson, L., Scheinin, N.M., Parkkola, R., Saunavaara, J., Lähdesmäki, T., Lehtola, S.J., Keskinen, M., Lewis, J.D., Evans, A.C., Karlsson, H., 2019. Test-retest reliability of Diffusion Tensor Imaging metrics in neonates. Neuroimage 197. https://doi.org/10.1016/j.neuroimage.2019.04.067

Miller, S.P., Vigneron, D.B., Henry, R.G., Bohland, M.A., Ceppi-cozzio, C., Hoffman, C., Newton, N., Partridge, J.C., Ferriero, D.M., Barkovich, A.J., 2002. Serial Quantitative Diffusion Tensor MRI of the Premature Brain□: Development in Newborns With and Without Injury. J. Magn. Reson. Imaging 16, 621–632. https://doi.org/10.1002/jmri.10205

Mukherjee, P., Miller, J.H., Shimony, J.S., Conturo, T.E., Lee, B.C.P., Almli, C.R., Mckinstry, R.C., 2001. Normal Brain Maturation during Childhood□: Developmental Trends Characterized with. Radiology 221, 349–358.

Neil, J., Miller, J., Mukherjee, P., Hu, P.S., 2002. Diffusion tensor imaging of normal and injured developing human brain ± a technical review. NMR Biomed. 15, 543–552. https://doi.org/10.1002/nbm.784

Neil, J.J., Shiran, S.I., McKinstry, R.C., Schefft, G.L., Snyder, A.Z., Almli, C.R., Akbudak, E., Aronovitz, J.A., Miller, J.P., Lee, B.C., Conturo, T.E., 1998. Normal brain in human newborns: apparent diffusion coefficient and diffusion anisotropy measured by using diffusion tensor MR imaging. Radiology 209, 57–66.

Ni, H., Kavcic, V., Zhu, T., Ekholm, S., Zhong, J., 2006. Effects of Number of Diffusion Gradient Directions. Am.J.Neuradiol 27, 1776–1781.

Oguz, I., Farzinfar, M., Matsui, J., Budin, F., Liu, Z., Gerig, G., Johnson, H.J., Styner, M., 2014. DTIPrep: Quality control of diffusion-weighted images. Front Neuroinform. 8, 1–11. https://doi.org/10.3389/fninf.2014.00004

Oishi, K., Mori, S., Donohue, P.K., Ernst, T., Anderson, L., Buchthal, S., Faria, A., Jiang, H., Li, X., Miller, M.I., van Zijl, P.C.M., Chang, L., 2011. Multi-contrast human neonatal brain atlas: Application to normal neonate development analysis. Neuroimage 56, 8–20. https://doi.org/10.1016/j.neuroimage.2011.01.051

Papadakis, N.G., Murrills, C.D., Hall, L.D., Huang, C.L.H., Adrian Carpenter, T., 2000. Minimal gradient encoding for robust estimation of diffusion anisotropy. Magn. Reson. Imaging 18, 671–679. https://doi.org/10.1016/S0730-725X(00)00151-X

Partridge, S.C., Mukherjee, P., Henry, R.G., Miller, S.P., Berman, J.I., Jin, H., Lu, Y., Glenn, O.A., Ferriero, D.M., Barkovich, A.J., Vigneron, D.B., 2004. Diffusion tensor imaging□: serial quantitation of white matter tract maturity in premature newborns. Neuroimage 22, 1302–1314. https://doi.org/10.1016/j.neuroimage.2004.02.038

Pierpaoli, C., Basser, P.J., 1996. Toward a quantitative assessment of diffusion anisotropy. Magn. Reson. Med. 36, 893–906. https://doi.org/10.1002/mrm.1910360612

Prayer, D., Prayer, L, 2003. Diffusion-weighted magnetic resonance imaging of cerebral white matter de v elopment Eur. J. Radiol. 45, 235–243.

Righini, A., Bianchini, E., Parazzini, C., Gementi, P., Ramenghi, L., Baldoli, C., Nicolini, U., Mosca, F., Triulzi, F., 2003. Apparent Diffusion Coefficient Determination in Normal Fetal Brain□: A Prenatal MR Imaging Study. AJNR. Am. J. Neuroradiol. 24, 799–804.

Sadeghi, N., Prastawa, M., Fletcher, P.T., Vachet, C., Wang, B., Gilmore, J., Gerig, G., 2013. Multivariate modeling of longitudinal MRI in early brain development with confidence measures, in: IEEE 10th International Symposium on Biomedical Imaging (ISBI). IEEE, pp. 1400–1403.

Sakaie, K., Lee, J., Debbins, J.P., Arnold, D.L., Melhem, E.R., Philips, M.D., Lowe, M., 2012. A Validation Study of Multicenter Diffusion Tensor Imaging□: Reliability of Fractional Anisotropy and Diffusivity Values. AJNR Am.J.Neuroradiol. 33, 695–700.

Skare, S., Hedehus, M., Moseley, M.E., Li, T.Q., 2000. Condition Number as a Measure of Noise Performance of Diffusion Tensor Data Acquisition Schemes with MRI. J. Magn. Reson. 147, 340–352. https://doi.org/10.1006/jmre.2000.2209

Smith, S.M., Jenkinson, M., Johansen-Berg, H., Rueckert, D., Nichols, T.E., Mackay, C.E., Watkins, K.E., Ciccarelli, O., Cader, M.Z., Matthews, P.M., Behrens, T.E.J., 2006. Tract-based spatial statistics: voxelwise analysis of multi-subject diffusion data. Neuroimage 31, 1487–505. https://doi.org/10.1016/j.neuroimage.2006.02.024

Tamnes, C.K., Roalf, D.R., Goddings, A.L., Lebel, C., 2018. Diffusion MRI of white matter microstructure development in childhood and adolescence: Methods, challenges and progress. Dev. Cogn. Neurosci. 33, 161–175. https://doi.org/10.1016/j.dcn.2017.12.002

Tanner, S.F., Ramenghi, L.A., Ridgway, J.P., Berry, E., Saysell, M.A., Martinez, D., Arthur, R.J., Smith, M.A., Levene, M.I., 2000. Quantitative Comparison of Intrabrain Diffusion in Adults and Preterm and Term Neonates and Infants. Am. J. Roentgenol. 174, 1643–1649.

Team, R.C., 2020. R: A Language and Environment for Statistical Computing [WWW Document]. URL https://www.r-project.org

Vaessen, M.J., Hofman, P.A.M., Tijssen, H.N., Aldenkamp, A.P., Jansen, J.F.A., Backes, W.H., 2010. The effect and reproducibility of different clinical DTI gradient sets on small world brain connectivity measures. Neuroimage 51, 1106–1116.https://doi.org/10.1016/j.neuroimage.2010.03.011

Wang, Y.J., Abdi, H., Bakhadirov, K., Diaz-arrastia, R., Devous, M.D.S., 2012. A comprehensive reliability assessment of quantitative diffusion tensor tractography. Neuroimage 60, 1127–1138. https://doi.org/10.1016/j.neuroimage.2011.12.062

Widjaja, E., Mahmoodabadi, S.Z., Rea, D., Moineddin, R., Vidarsson, L., Nilsson, D., 2009. Effects of gradient encoding and number of signal averages on fractional anisotropy and fiber density index in vivo at 1.5 tesla. Acta radiol. 50, 106–113. https://doi.org/10.1080/02841850802555646

Winkler, A.M., Ridgway, G.R., Webster, M.A., Smith, S.M., Nichols, T.E., 2014. Permutation inference for the general linear model. Neuroimage 92, 381–397. https://doi.org/10.1016/j.neuroimage.2014.01.060

