## supplemental material for "Effect of number of diffusion encoding directions in Neonatal Diffusion Tensor Imaging using Tract-Based Spatial Statistical analysis"

**Supplementary Information Table 1** Publications involving Diffusion Weighted Imaging (DWI) and Diffusion Tensor Imaging (DTI) in study of brain development *in vivo*, including infants of less than 24 months after birth, with number of tensor directions, ages of scanned subjects and analysis type.

| **Study** | **Field Strength** | **Main Diffusion parameters** | **Analysis method** | **Diffusion Data Quality Assurance** | **Infants Included** |
| --- | --- | --- | --- | --- | --- |
| **Neil et al. 1998** | **1.5T** | **4 tetrahedron directions with b=800, 3 additional orthogonal with b=340** | **T2-based regions of interest** | **Manual motion detection, discarding cases with bad directions.** | **12 term, 10 preterm, 11 male, 11 female** |
| **Mukherjee at al. 2001** | **1.5T** | **4 tetrahedron directions with b=800, 3 additional orthogonal with b=338** | **DWI –based ROI with ANALYZEAVW software, 4 WM ROIs** | **Two-dimension aligment for motion and eddy current** | **1 day – 11 years, N=152** |
| **Forbes et al. 2002** | **1.5T** | **3 orthogonal directions with b=1000** | **4 GM, 4 WM average ADC map based ROIs** | **N/A** | **0 - 12 months N=40, 19 male, 21 female** |
| **Miller at al. 2002** | **1.5T** | **6 orthogonal directions with b=600** | **9 ADC map based ROIs** | **In-house software for rotationally invariant parameter maps** | **Premature, GA < 36 weeks, N=23** |
| **McGraw et al. 2002** | **1.5T** | **6 orthogonal directions with b=1000** | **8 FA map based ROIs with Functool software, in Corpus Callosum, Internal Capsule, Cerebral Peducle** | **N/A** | **4 days - 12 months (Group 1), N=40** |
| **Righini et al. 2003** | **1.5T** | **3 orthogonal directions with b=600** | **T2-based ROIs with Functool software in basal ganglia, frontal white matter, occipital white matter, CSF** | **Quality control during scan for individual directions, selection of better motion artifact-free for ROI drawing** | ***in utero*, GA 22 - 35 weeks, N=15** |
| **Partridge et al. 2004** | **1.5T** | **6 orthogonal directions with b=600** | **FA and MD map based delineations of 12 ROIs in WM** | **Measures taken in acquisition to allow sleep and minimize motion artefacts,** | **Preterm, GA 28 - 39 weeks, N=14** |
| **Maas et al. 2004** | **1.5T** | **6 orthogonal directions with b=600** | **FA, ACD and direction of main eigenvector** | **Visual inspection** | **Preterm, GA 24 and 25, N=2** |
| **Berman et al. 2005** | **1.5T** | **6 directions with b=600** | **Manual seed ROI placement, tracktography ROI analysis** | **Non-linear co-regitration for motion correction, manual quality assurance for ROIs** | **GA 28-43 weeks, N=27** |
| **Yoo et al. 2005** | **1.5T** | **6 directions with b=1000** | **Manual seed ROI placement, tracktography ROI analysis** | **Manual screening of DTI image quality** | **GA 28-40 weeks, N=6** |
| **Bui et al. 2006** | **1.5T** | **6 orthogonal directions with b=700** | **3 DWI, T1, T2 -based ROIs in WM, with in-house software** | **DTI studies were excluded if significant motion or technical artefacts found** | ***in utero*, GA 31-37 weeks, N=24** |
| **Hermoye et al. 2006** | **1.5T** | **32 directions with b=700** | **12 FA map based ROIs in WM tracts** | **Anesthesia to avoid motion artefacts with patients, healthy scanned while in sleep** | **0-54 months, N=30, 7 healthy, 23 patients, 17 boyt 13 girls** |
| **Dudink et al. 2007** | **1.5T** | **25 directions with b=1000** | **FA-based manual ROIs in 4 WM tracts, bilateral** | **Preterm infants with severe motion excluded** | **Preterm 0-4 days (GA 25-32 weeks), N=28** |
| **Kasprian et al. 2008** | **1.5T** | **32 directions with b=700** | **Tract-based ROIs with FACT algorithm** | **Restricted head motion, detection of corrupted slices, co-registration, repeated analysis** | ***in utero*, GA 18-37 weeks, N=40** |
| **Jiang et al. 2009** | **1.5T** | **15 directions with b=500** | **Semi-automatic head segmentation. Co-registration based ROIs on DWI with manual refinements.** | **Slices with in-plane motion detectied manually and exclusion** | ***in utero*, 24 < GA < 34, N=8** |
| **Aeby et al. 2009** | **1.5T** | **32 directions with b=600** | **Template-baced ROI placement** | **Visual inspection for motion artefacts, directions with artefacts excluded. Visual inspection of co-registration and FA, MD maps.** | **24<GA<41 healthy preterm N=22 healthy term N=6** |
| **Aeby et al. 2012** | **1.5T** | **32 directions with b=600** | **Template-baced ROI placement** | **N/A** | **65 preterm, 35 boys, 30 girls** |
| **Geng et al. 2012** | **3T** | **6 directions with b=1000** | **Template-based nad tracktography-based ROIs** | **Screening for motion artifacts, missing and corrupted sections with DTIprep** | **2-4 weeks, 63 neonates** |
| **Sadeghi et al. 2013** | **N/A** | **N/A** | **Template-based ROIs using T2** | **N/A** | **approximately 2 weeks, 1 year and 2 years, 26 healthy subjects** |
| **Friedrichs-Maeder et al. 2017** | **3T** | **30 directions with b=1000** | **ADC with connectomics** | **N/A** | **Preterm, N=9, 4 male, 5 female** |
| **Lebenberg at al. 2019** | **3T** | **30 directions with b=700** | **FA and AD** | **Visual inspection** | **Maturation age 3-21 weeks, N=17, 10 male, 7 female** |
| **Batalle et al. 2019** | **3T** | **64 with b=2500** | **ROI-based FA, MD** | **Visual inspection. Sedation offered in acquisition. Images with motion artefacts excluded from study.** | **Preterm GA 25-47 weeks, N=99, 60 male, 39 female** |

**Supplemental Information Figure 1**

Mean MD, AD, RD skeleton rendering on 133 neonatal subjects with 54 diffusion encoding directions

Mean Diffusivity

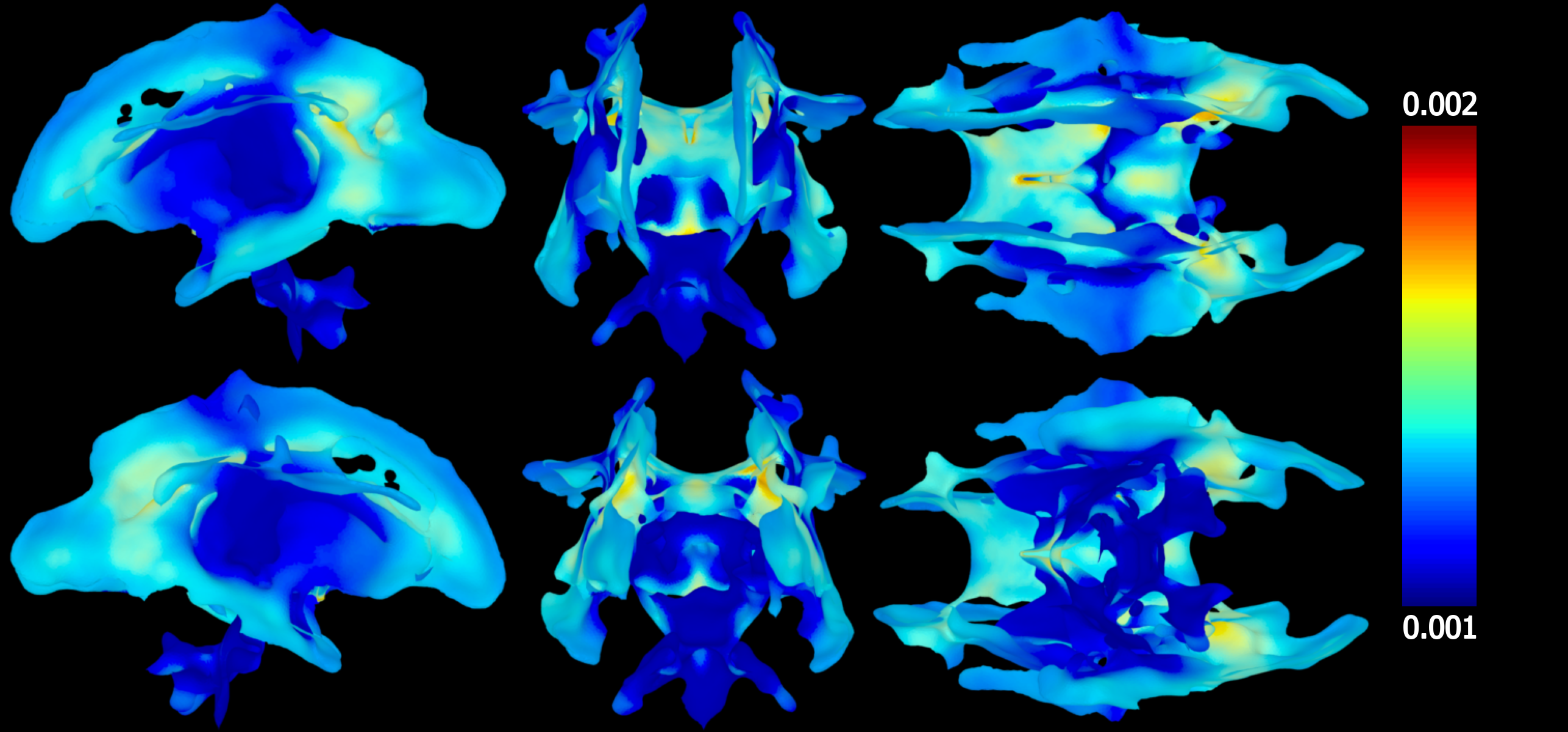

Axial Diffusivity

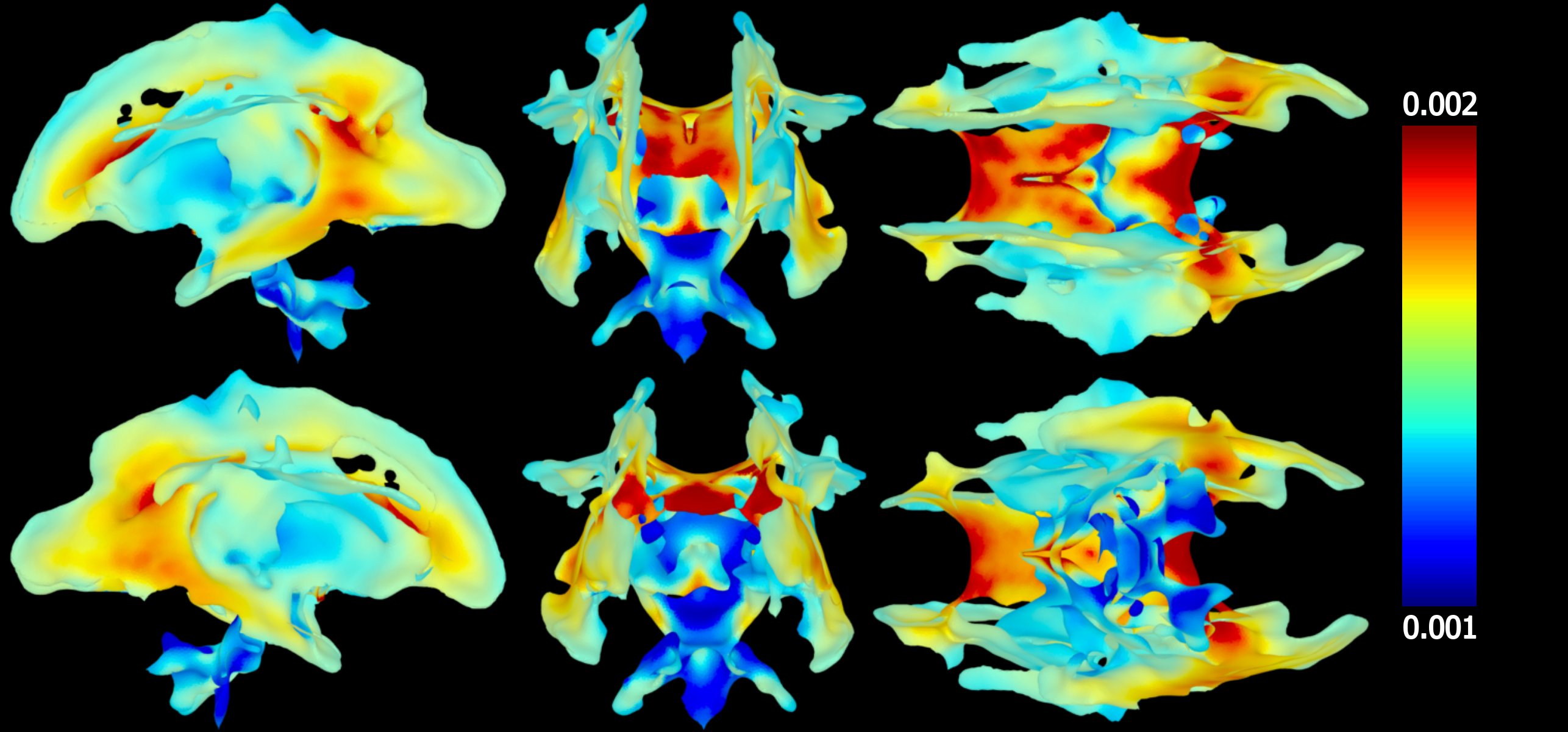

Radial Diffusivity

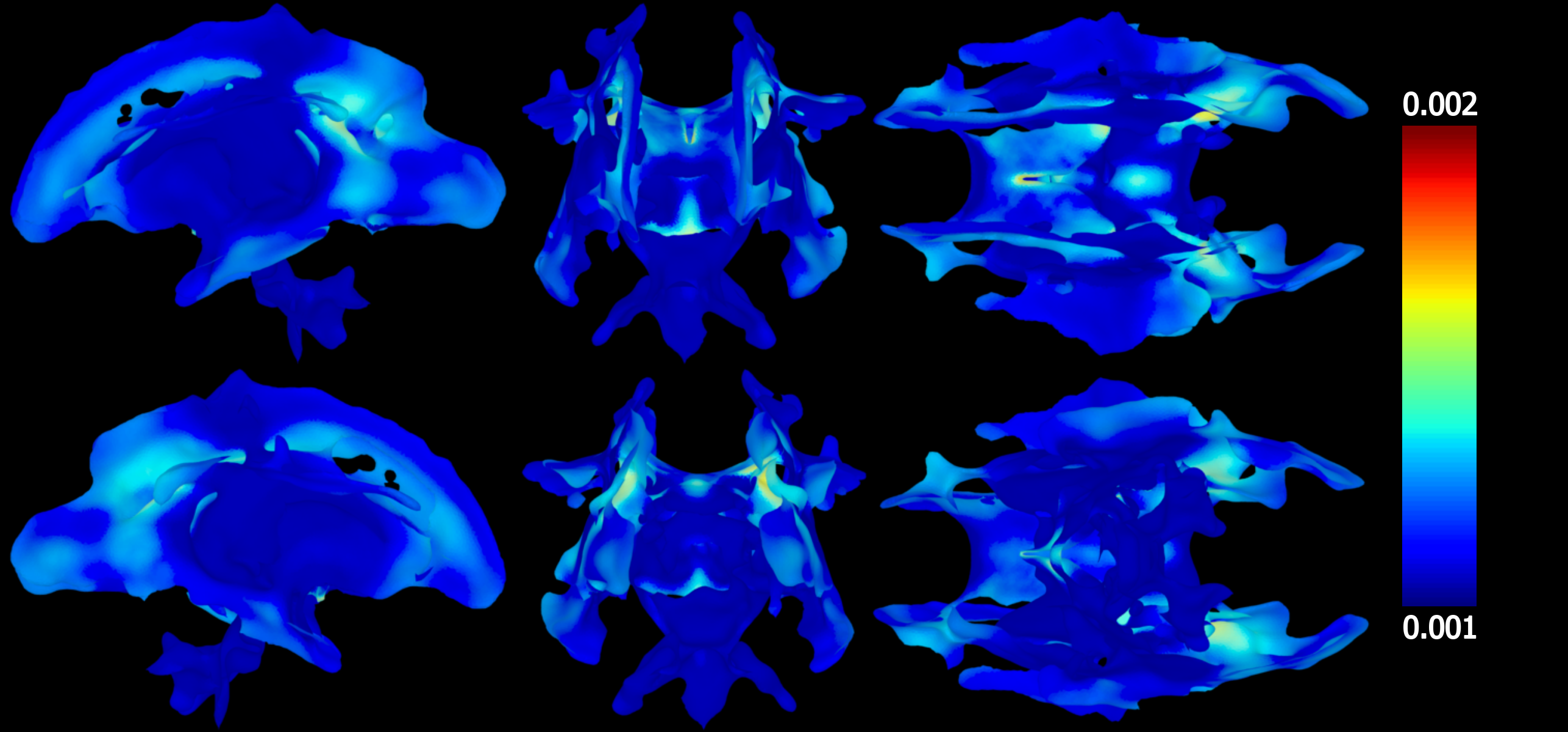

**Supplemental Information Figure 2**

Proportions of neonatal subjects (%) from 133 neonatal subjects, where difference in DTI scalar was larger than 10%. Comparison is from 6 (N6) to 48 (N48) diffusion encoding directions to reference values with 54 directions.

Mean Diffusivity

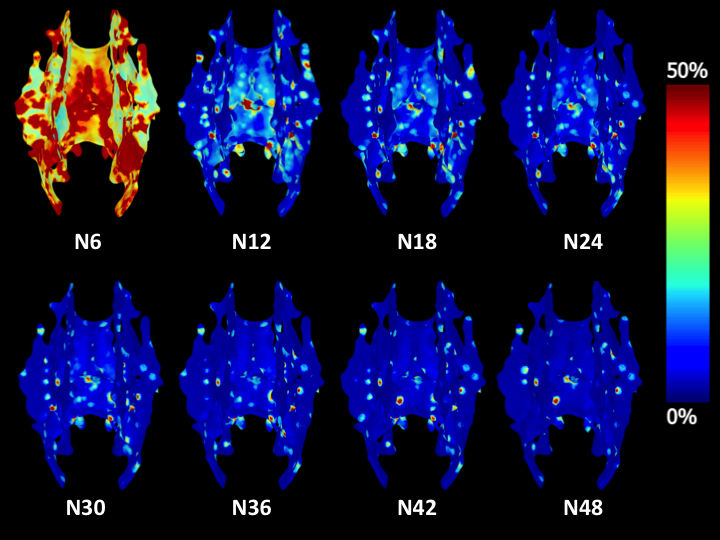

Axial Diffusivity

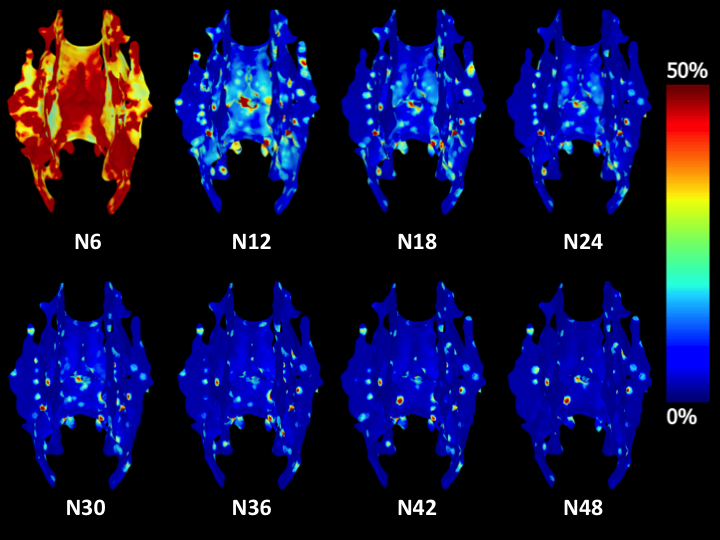

Radial Diffusivity

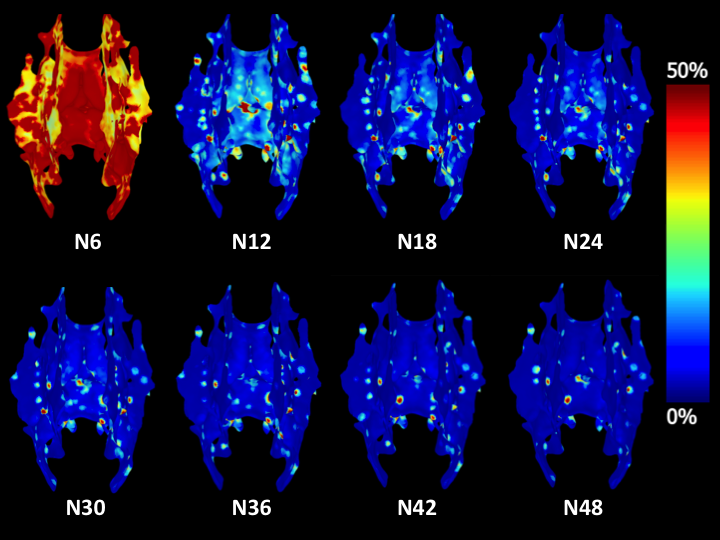

**Supplemental Information Figure 3**

Box plots of 6, 24 and 54 in main regions of JHU atlas for DTI scalars of FA, AD, MD and RD in 133 neonatal subjects.

Fractional Anisotropy

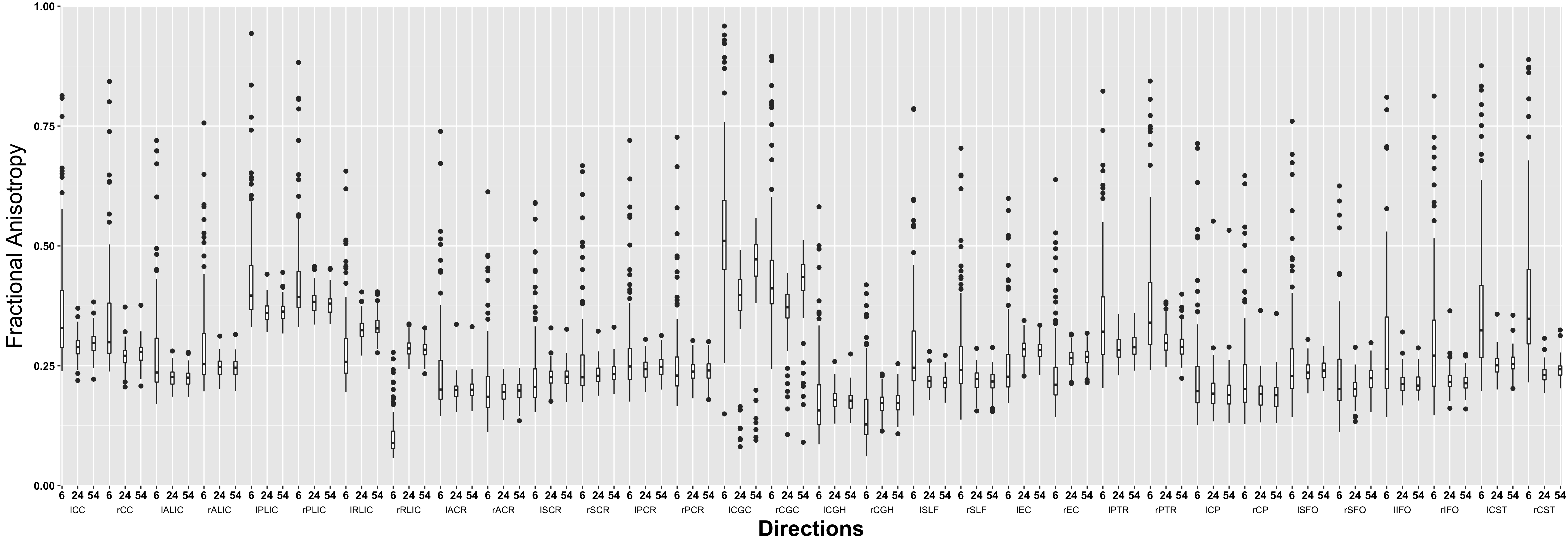

Axial Diffusivity

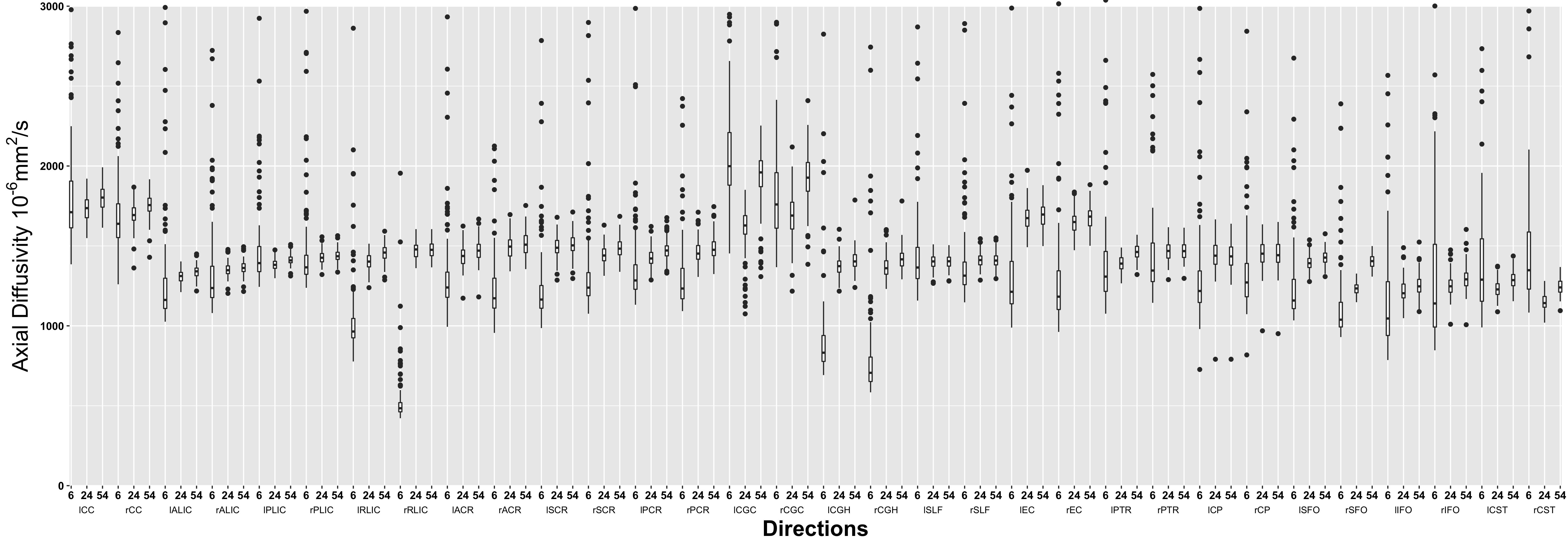

Mean Diffusivity

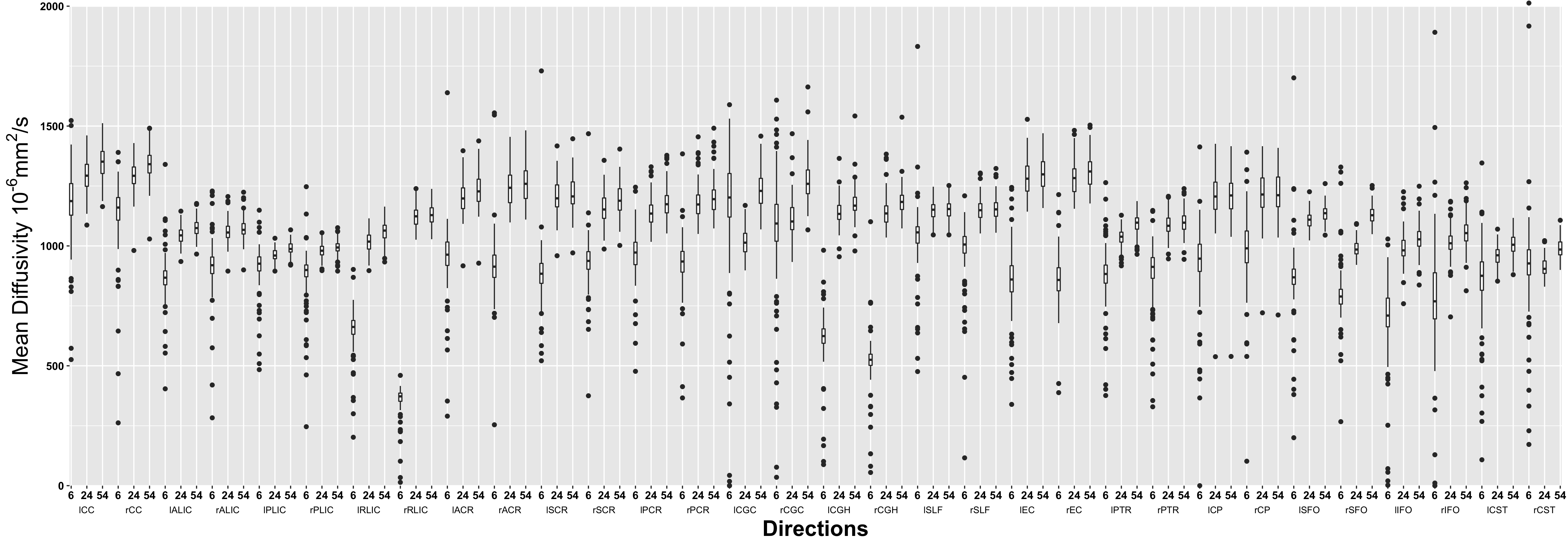

Radial Diffusivity

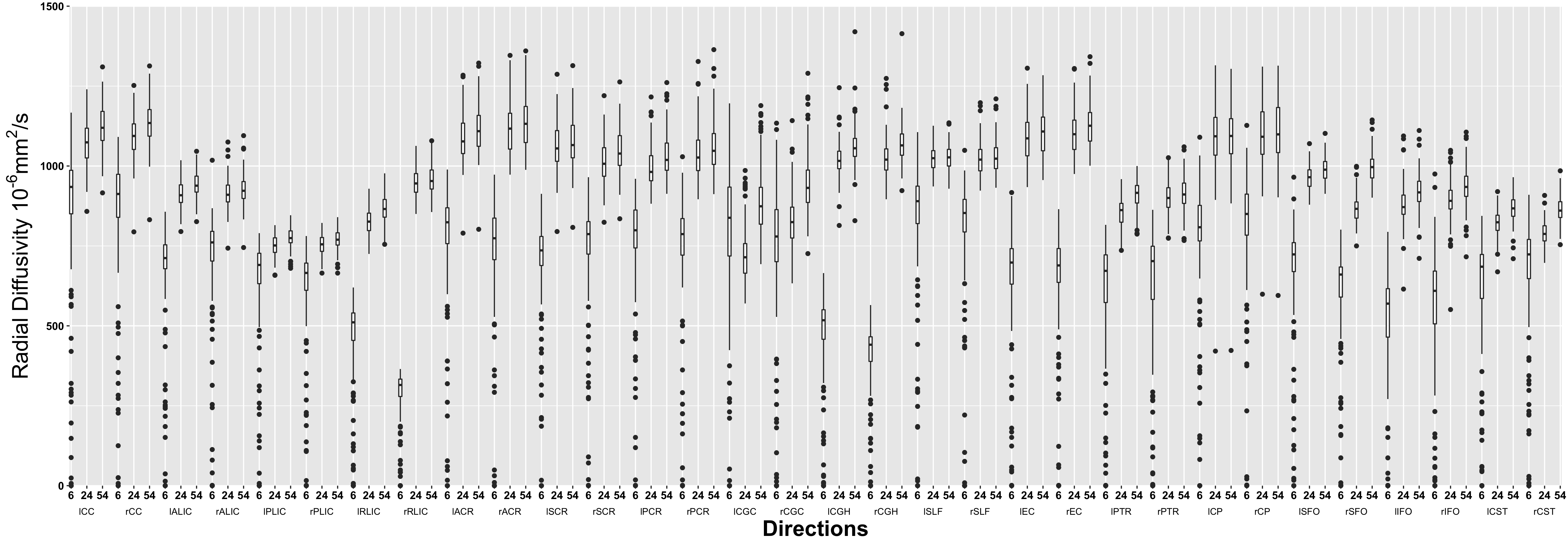

**Supplemental Information Figure 4**

Difference of MD, AD, RD (x10^-3^ mm^2^/s) between 54 and 24 directions in 133 neonatal subjects.

Mean Diffusivity

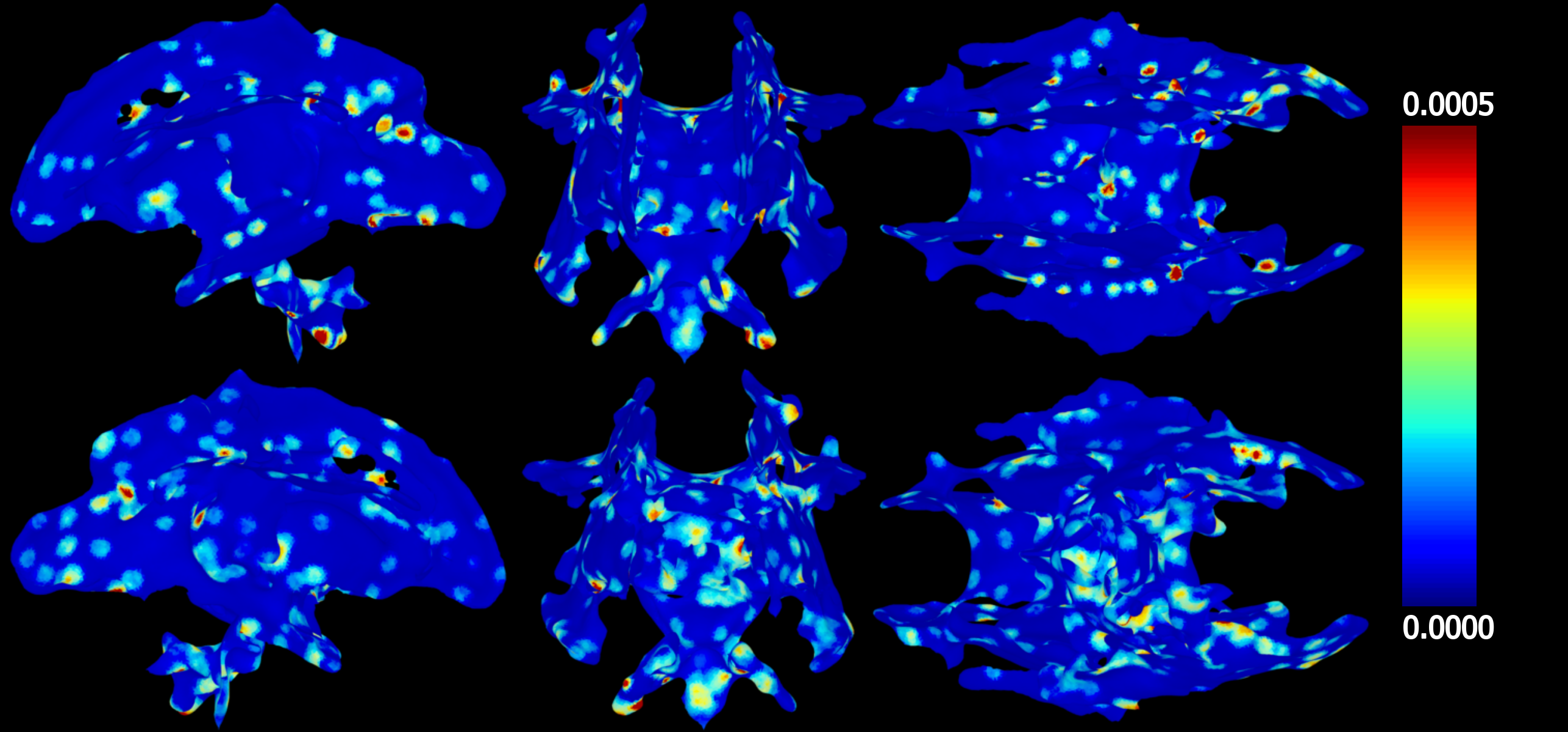

Axial Diffusivity

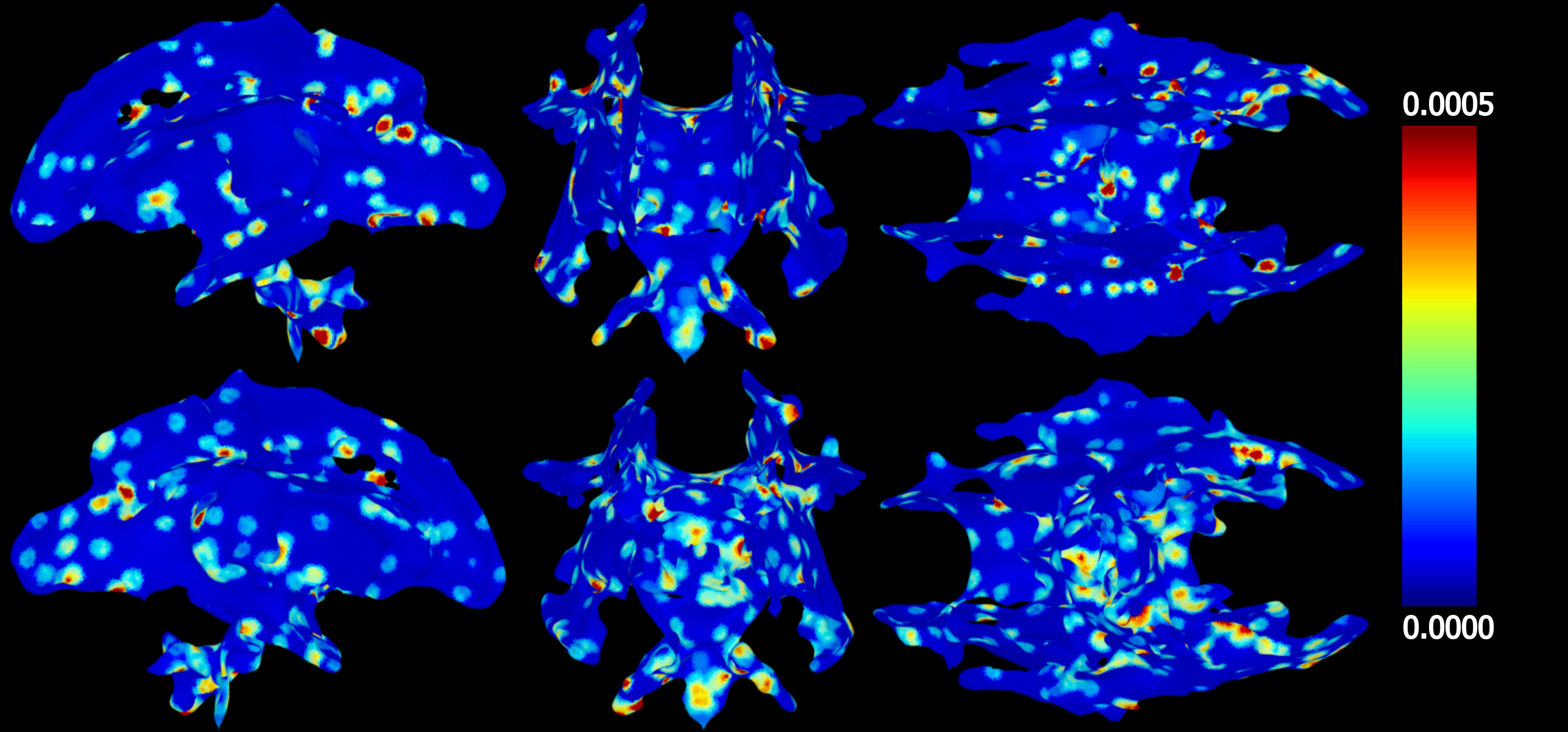

¨

Radial Diffusivity

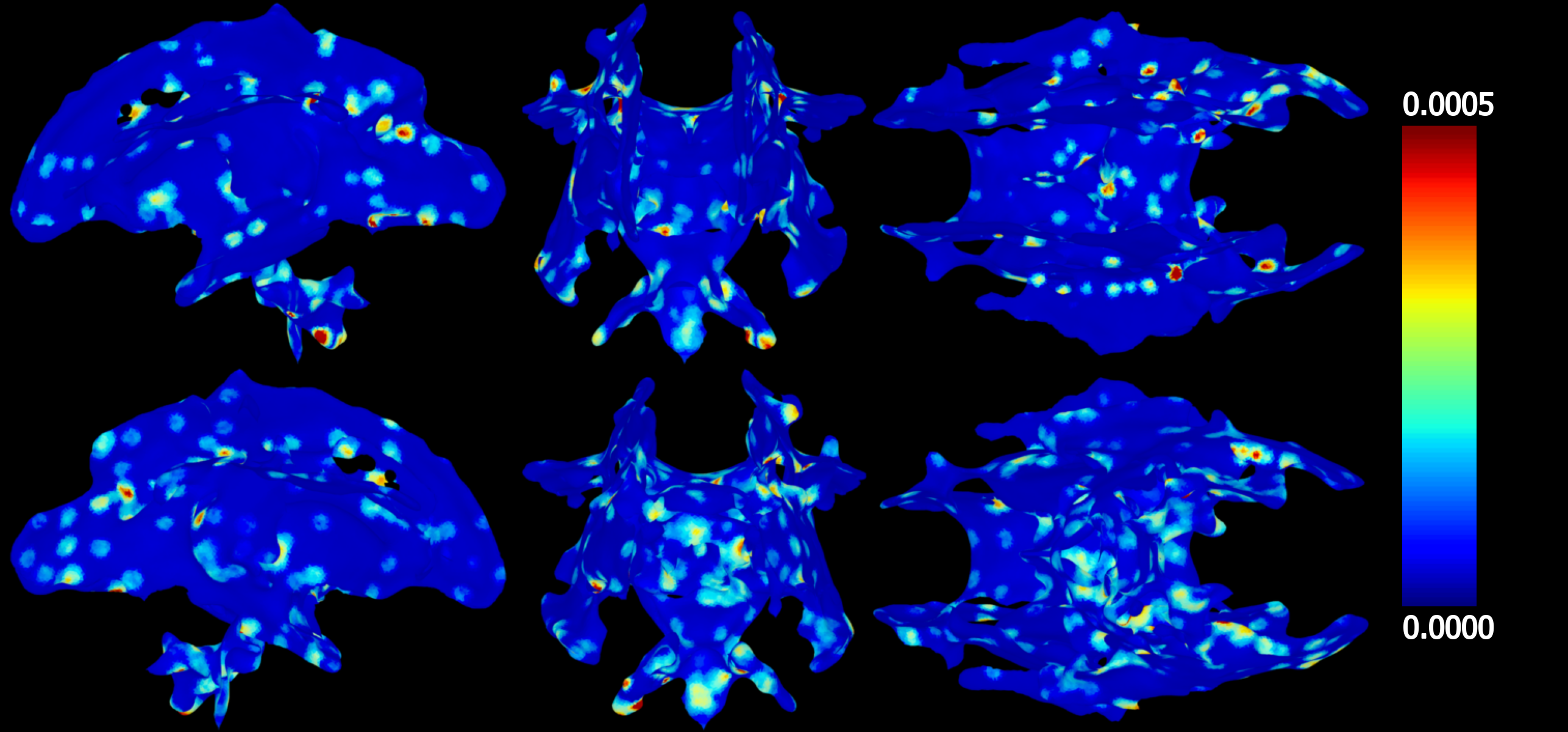

**Supplemental Information Figure 5**

ICC(2,1) of MD, AD, RD between 24, 30, 36, 42, 48 and 54 directions in 133 neonatal subjects.

Mean Diffusivity

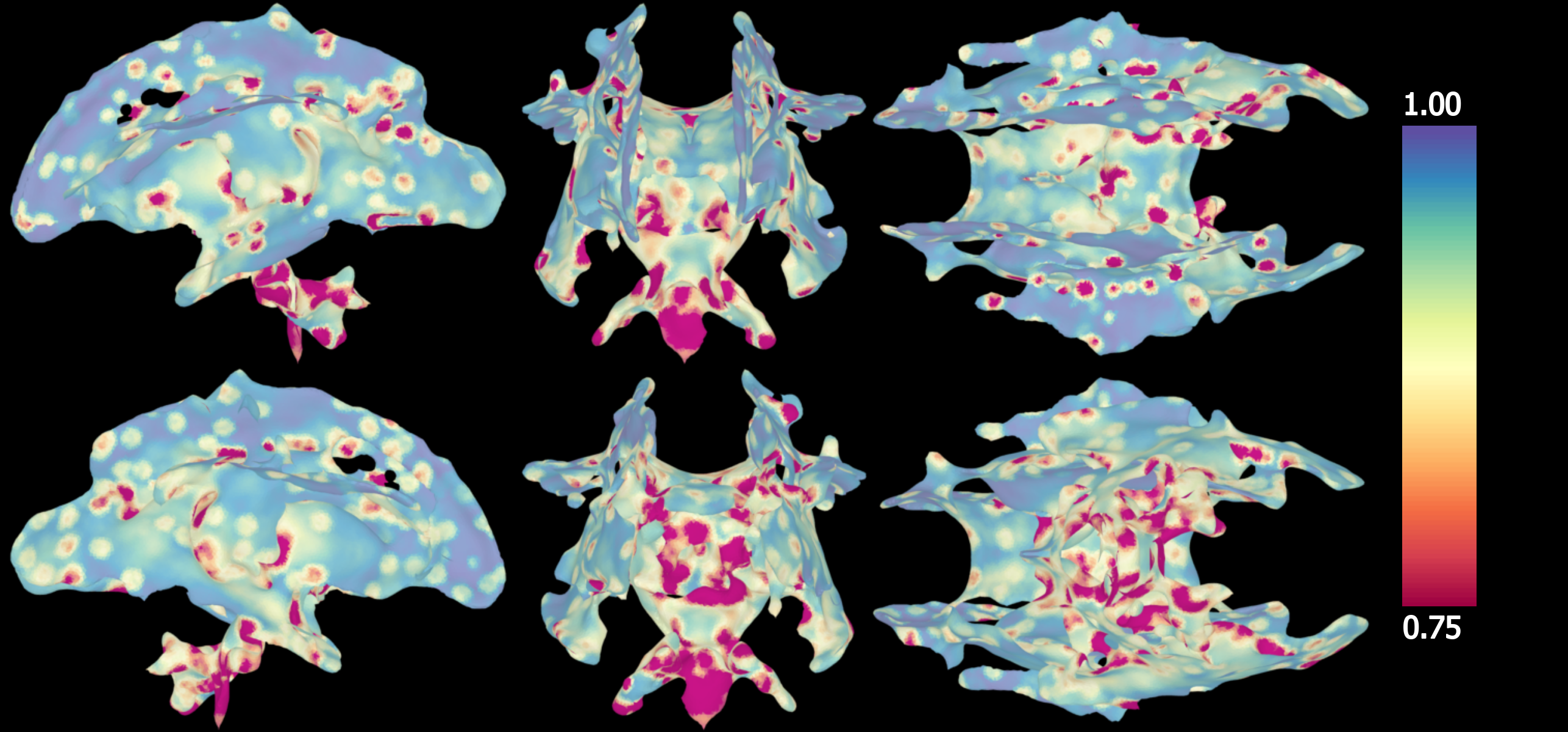

Axial Diffusivity

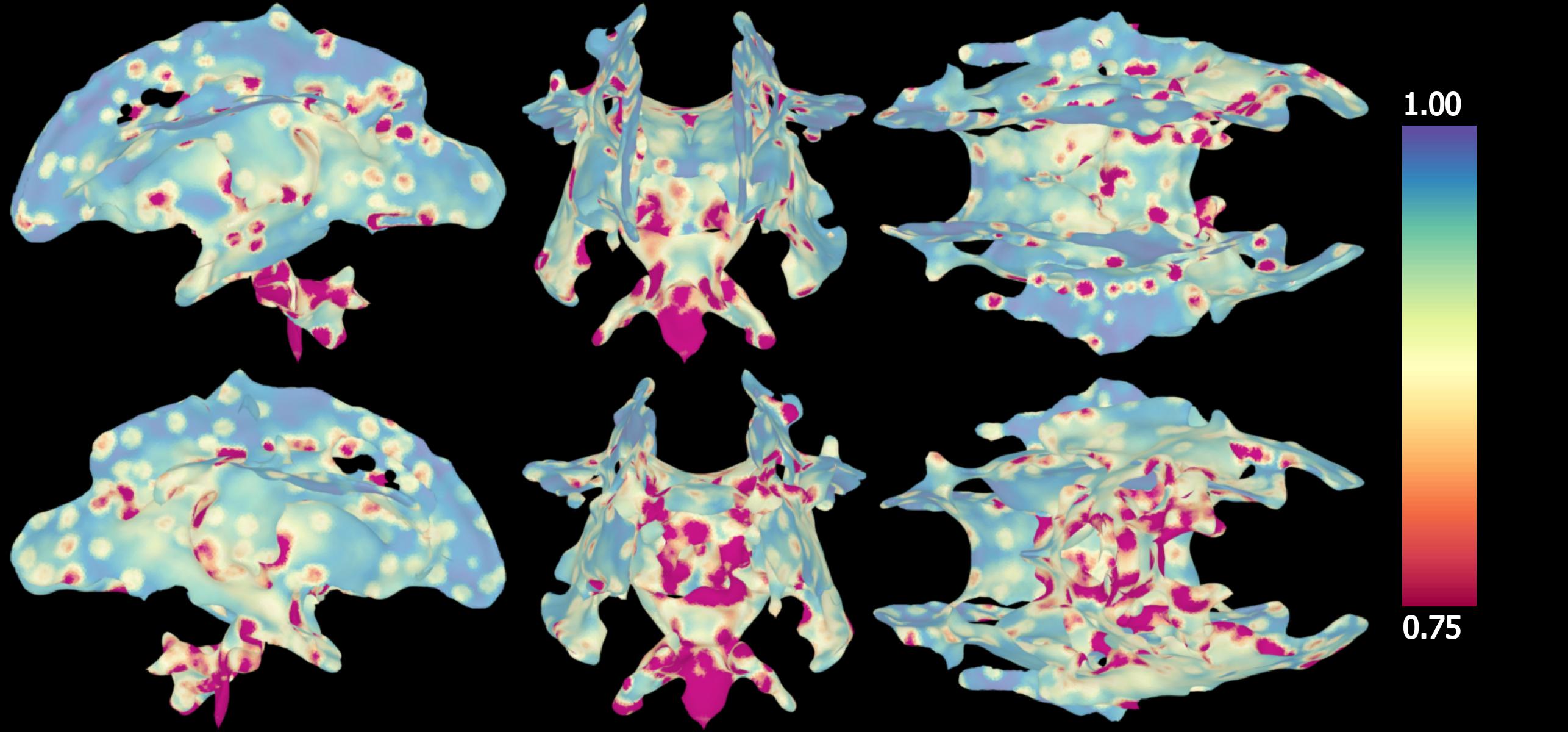

Radial Diffusivity

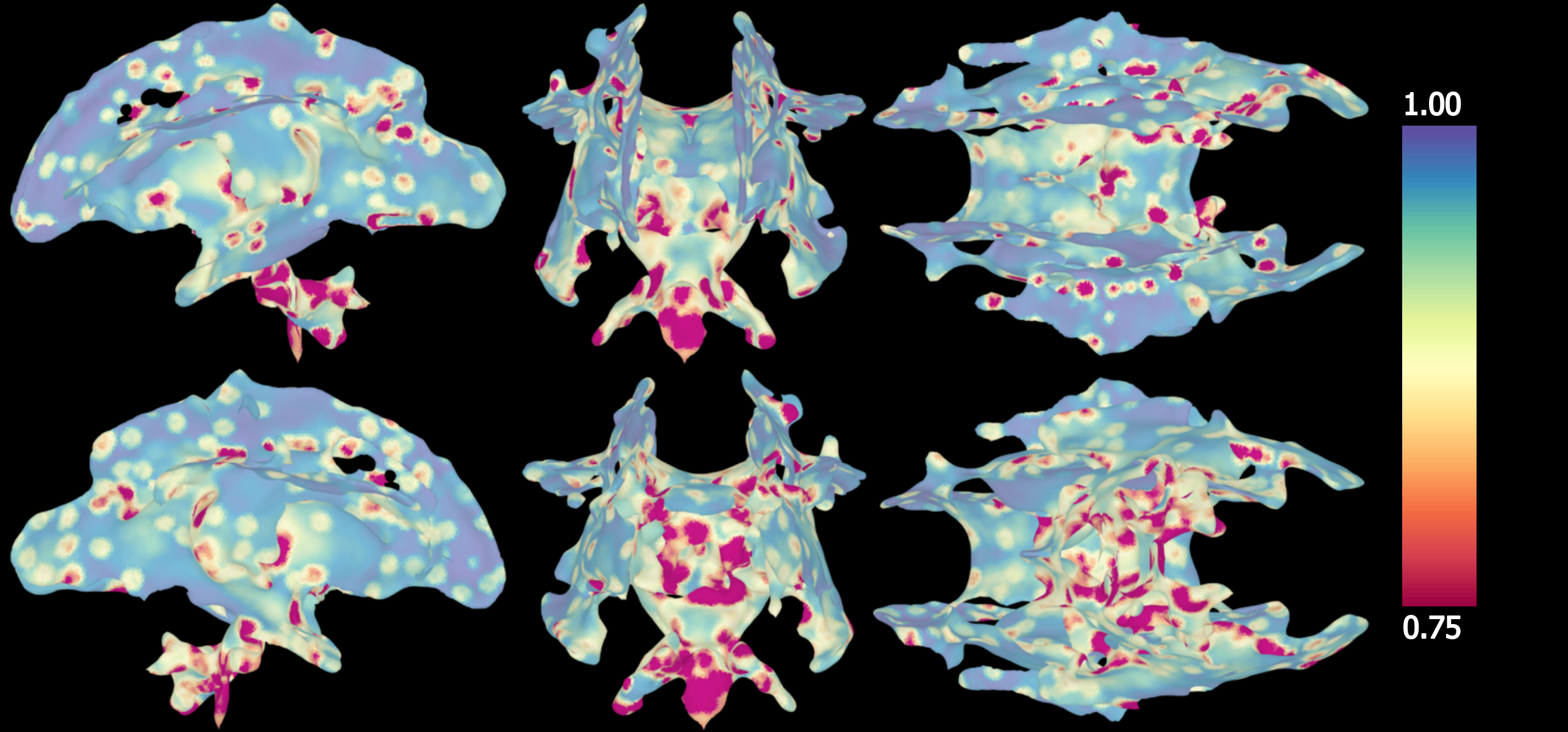
